# Regeneration of dopaminergic neurons in adult zebrafish depends on immune system activation and differs for distinct populations

**DOI:** 10.1101/367151

**Authors:** Lindsey J. Caldwell, Nick O. Davies, Leonardo Cavone, Karolina S. Mysiak, Svetlana A. Semenova, Pertti Panula, J. Douglas Armstrong, Catherina G. Becker, Thomas Becker

## Abstract

Adult zebrafish regenerate neurons in their brain, but the extent and variability of this capacity is unclear. Here we ask whether loss of various dopaminergic neuron populations is sufficient to trigger their functional regeneration. Genetic lineage tracing shows that specific diencephalic ependymo-radial glial progenitor cells (ERGs) give rise to new dopaminergic (Th^+^) neurons. Ablation elicits an immune response, increased proliferation of ERGs and increased addition of new Th^+^ neurons in populations that constitutively add new neurons, e.g. diencephalic population 5/6. Inhibiting the immune response attenuates neurogenesis to control levels. Boosting the immune response enhances ERG proliferation, but not addition of Th^+^ neurons. In contrast, in populations in which constitutive neurogenesis is undetectable, e.g. the posterior tuberculum and locus coeruleus, cell replacement and tissue integration are incomplete and transient. This is associated with loss of spinal Th^+^ axons, as well as permanent deficits in shoaling and reproductive behaviour. Hence, dopaminergic neuron populations in the adult zebrafish brain show vast differences in regenerative capacity that correlate with constitutive addition of neurons and depend on immune system activation.

## INTRODUCTION

The adult mammalian brain shows very limited neurogenesis after injury or neuronal loss, leading to permanent functional deficits ^1,2^. By contrast, the regenerative capacity of the CNS in adult zebrafish after injury is remarkable ^3-5^. However, relatively little is known about the capacity for regeneration and functional integration after loss of discrete cell populations in the fully differentiated adult CNS.

To study regeneration of distinct populations of neurons without physical damage, we ablated dopaminergic and noradrenergic neurons using 6-hydroxydopamine (6OHDA), which selectively ablates these neurons across vertebrates ^6-9^. In adult zebrafish, the dopaminergic system is highly differentiated. There are 17 distinct dopaminergic and noradrenergic brain nuclei, identified by immunohistochemistry for cytoplasmic Tyrosine hydroxylase (Th) and the related Th2, rate-limiting enzymes in dopamine and noradrenaline synthesis ^10,11^. Projections of Th^+^ brain nuclei are far-reaching, including long dopaminergic projections to the spinal cord from population 12 in the diencephalon and noradrenergic projections from the locus coeruleus (LC) in the brainstem. These projections are the only Th^+^ input to the spinal cord ^10,12,14^.

Functionally, dopamine, especially from the diencephalo-spinal projection from population 12, has roles in maturation and initiation of motor patterns in developing zebrafish ^15-18^. In addition, dopamine has been linked to anxiety-like behaviour in zebrafish ^19,20^. Dopaminergic neurons are constantly generated in the adult diencephalon ^21^, but it is unclear which populations receive new neurons and how this may change after ablation.

For regeneration of neurons to occur, ependymo-radial glia (ERG) progenitor cells need to be activated. ERGs have a soma that forms part of the ependyma and radial processes that span the entire thickness of the brain. After a CNS injury, these cells are either activated from quiescence or increase their activity in constitutively active adult proliferation zones to regenerate lost neurons ^3,5,22^. Activation could occur via damage to the highly branched ERG processes or early injury signals. Remarkably, the microglial/macrophage reaction following a mechanical lesion has been shown to be both necessary and sufficient for regenerative proliferation of ERGs and neurogenesis in the adult zebrafish telencephalon ^23^. The immune response also promotes neuronal regeneration in the spinal cord of larval zebrafish after a lesion ^24^. Hence, it might also play a role in the regenerative response after discrete neuronal loss without injury.

We find that locally projecting dopaminergic neurons in the diencephalon are regenerated from specific ERGs, whereas large Th^+^ neurons with spinal projections are only transiently replaced, associated with permanent and specific functional deficits in shoaling and mating behaviour. Inhibiting the immune response abolished ablation-induced regeneration. Hence, we demonstrate an unexpected heterogeneity in regenerative capacity of functionally important dopaminergic neurons in the adult zebrafish and essential functions of the immune response.

## MATERIAL AND METHODS

### Animals

All fish were kept and bred in our laboratory fish facility according to standard methods ^25^, and all experiments had been approved by the British Home Office. We used wild type (*wik*) and *Tg(olig2:DsRed2)* ^26^, abbreviated as *olig2:DsRed; Tg(gfap:GFP)* ^27^, abbreviated as *gfap:GFP;Tg(slc6a3:EGFP)* ^28^, abbreviated as *dat:GFP,* and *Tg(*her4.1*:TETA-GBD-2A-mCherry)* ^29^, abbreviated as *her4.3*:mCherry, transgenic reporter lines. Note that zebrafish nomenclature treats *her4.1* and *her4.3* as synonymous (https://zfin.org/ZDB-TGCONSTRCT-110825-6). For genetic lineage tracing, we used *Tg(-3her4.3:Cre-ERT2)* ^30^ crossed with *Tg(actb2:LOXP-mCherry-LOXP-EGFP)* ^31^, as previously described ^32^. Adult (> 4 months of age) male and female fish were used for the experiments.

### Bath application of substances

For dexamethasone treatment, fish were immersed in 15 mg/L dexamethasone (Sigma-Aldrich, D1756) or vehicle (DMSO) in system water. For lineage tracing experiments, fish were immersed in 1 μM 4-hydroxytamoxifen (Sigma-Aldrich, H6278) in system water with tanks protected from light. Fish were transferred into fresh drug/vehicle every other day.

### Intraventricular injections

Fish were anaesthetised in MS222 (Sigma-Aldrich,1:5000 % w/v in PBS) and mounted in a wet sponge to inject the third ventricle from a dorsal approach using a glass capillary, mounted on a micromanipulator. Using sharp forceps, a hole was made into the skull covering the optic tectum and the needle was advanced at a 45° angle from the caudal edge of the tectum into the third ventricle. The capillary was filled with a 10 mM solution of 6OHDA (6-Hydroxydopamine hydrobromide, Sigma-Aldrich, product number: H116) in H_2_O and 0.12% of a fluorescent dextran-conjugate (Life Technologies, product number: D34682) to ablate Th^+^ cells, or with fluorescently labelled Zymosan A (from *Saccaromyces cerevisiae)* bioparticles at a concentration of 10 mg/mL (Life Technologies, product number: Z23373) to stimulate the microglial response. LTC4 (Cayman Chemicals, product number: 20210) was injected at a concentration of 500 ng/ml in 0.45% ethanol in H_2_O. Sham-injected controls were generated by injecting vehicle solutions.

A pressure injector (IM-300 microinjector, Narishige International, Inc. USA) was used to inject 0.5 to 1.0 μL of the solution. Distribution of the solution throughout the ventricular system was verified under a fluorescence-equipped stereo-microscope. This injection technique only induced a localised microglia reaction surrounding the point where the capillary penetrated the optic tectum, but not close to any of the Th^+^ populations of interest.

### Intraperitoneal injections

Fish were anaesthetised in MS222 and injected on a cooled surface on their left side with a 30½ G needle. Per application, 25 μl of 16.3 mM EdU (Invitrogen) was injected intraperitoneally. EdU was dissolved in 15% DMSO and 30% Danieau's solution in distilled water.

Haloperidol (Sigma-Aldrich, product number: H1512) was injected at a volume of 25 μl and a concentration of 80 μg/ml in PBS for each injection. This roughly equates to 4 mg/kg, twice the concentration shown to be effective in salamanders ^8^.

### Quantitative RT-PCR

Brains were dissected without any tissue fixation and sectioned on a vibrating-blade microtome. RNA was isolated from a horizontal section 200 μm thick at the level used for analysis of proliferating ERGs around the ventricle (refer to Fig. 5A for section level) using the RNeasy Mini Kit (Qiagen, 74106). cDNA synthesis was performed using the iScript™ cDNA Synthesis Kit (Bio-Rad, 1708891). Standard RT-PCR was performed using SsoAdvanced™ Universal SYBR^®^ Green Supermix (Bio-Rad, 172-5271). qRT-PCR (annealing temperature 58 °C) was performed using Roche Light Cycler 96 and relative mRNA levels were determined using the Roche Light Cycler 96 SW1 software. Samples were run in duplicates and expression levels were normalized to the level of 18S ribosomal RNA. Primers were designed to span an exon-exon junction using Primer-BLAST. Primer sequences:

TNF-α FW 5’-TCACGCTCCATAAGACCCAG-3’, RV 5’-

GATGTGCAAAGACACCTGGC-3’, il-1β FW 5’-

ATGGCGAACGTCATCCAAGA-3’, RV 5’-GAGACCCGCTGATCTCCTTG-3’,

18S FW 5’-TCGCTAGTTGGCATCGTTTATG-3’,RV 5’-

CGGAGGTTCGAAGACGATCA-3’.

### HPLC

Brains were dissected without any tissue fixation and frozen. HPLC analysis was performed as described ^33^.

### Immunohistochemistry

We used mouse monoclonal antibody 4C4 (1:50; HPC Cell Cultures, Salisbury, UK, catalogue number: 92092321) to label microglia. The antibody labels microglia in the brain, but not peripheral macrophages ^34^. We used a chicken antibody to green fluorescent protein (GFP) (1:500; Abcam, Cambridge, MA, USA, designation: ab13970); a mouse monoclonal antibody to the proliferating cell nuclear antigen (PCNA) (1:1000; Dako, Sigma-Aldrich, St Louis, MO, USA, designation: M0879); a mouse monoclonal antibody to tyrosine hydroxylase (Th) (1:1000; Merck Millipore, Billerica, MA, US, designation: MAB318). Suppliers for the appropriate fluorescence or biotin-labelled antibodies were Stratech Scientific, Sydney, Australia and Vector Laboratories, Burlingame, CA, USA, respectively. Dilutions of secondary antibodies followed the manufacturers' recommendations.

Immunofluorescent labelling of 50 μm sections was carried out as previously described ^35^. Briefly, brains from perfusion-fixed (4% paraformaldehyde) animals were dissected, sectioned on a vibrating-blade microtome, incubated with primary antibody at 4°C overnight, washed, incubated in secondary antibody for 45 min at room temperature, washed and mounted in glycerol. All washes were 3 times 15 minutes in PBSTx (0.1% Triton X 100 in PBS).

For colorimetric detection of Th, a biotinylated secondary antibody was used, followed by the ABC reaction using the Vectastain ABC kit (Vector Laboratories, Burlingame, USA) according to the manufacturer's recommendations. The colour was developed using diaminobenzidine solution (1:120 diaminobenzidine; 2 μl/ml of 1% stock NiCl_2_ and 2 μl/ml of 1% stock CoSO_4_ in PBS) pre-incubation (30 min at 4°C), followed by addition of 30% hydrogen peroxide. Sections were mounted, dried and counterstained in neutral red staining solution (4% acetate buffer (pH 4.8) and 1% neutral red in dH_2_O) for 6 min, followed by differentiation in 70% and 95% ethanol.

### EdU detection

To detect EdU, we used Click-iT^®^ EdU Alexa Fluor^®^ 488 or 647 Imaging Kits (Molecular Probes) according to the manufacturer's recommendations. Briefly, 50 μm sections from perfusion-fixed brains were incubated in Click-iT reaction buffer for three hours in the dark at room temperature, washed 3 ×10 min in 0.3% PBSTx and once in PBS. After that, sections were mounted in 70% glycerol or underwent immunofluorescent labelling as above.

### TUNEL labelling

TUNEL labelling was carried out as described ^36^ using the *in situ* TMR cell death detection kit (Roche) according to the manufacturer's recommendations. In brief, sections were incubated with reaction mix in the dark at 37°C for 60 min. This was followed by immunolabelling as described above.

### Quantification of cells and axons

All counts were carried out with the observer blinded to the experimental condition. For colorimetric immunohistochemistry of Th, cell profiles were counted for individual brain nuclei, identified by neutral red counterstain. Innervation density of labelled axons was semi-quantitatively determined by determining the average pixel brightness for a region of interest using Image J.

In fluorescently labelled sections, cells were stereologically counted in confocal image stacks, as described ^35^. Double-labelling of cells was always assessed in single optical sections (<2 μm thickness). Fluorescently labelled axons in the spinal cord were quantified using automatic functions in Image J as described ^14^.

### Behavioural tests

All behaviour tests, comparing between 6OHDA-injected and sham-injected animals, were performed when at least seven days had passed after injection. All recordings were made with a Sony ExwaveHAD B&W video camera and videos were analysed using Ethovision XT7 tracking software (Noldus, Leesberg, USA), except for shoaling analysis (see below).

For the open field test, fish swimming was recoded in a round tank (16.3 cm diameter, 8 cm water depth) for 6 min after 2 minutes acclimatization time. The software calculated the total distance moved and the average velocity of fish.

For the light/dark test, a tank (10 cm × 20 cm, 8 cm water depth) was illuminated from below with half of the area blocked from the light. The time spent in the illuminated area was recorded in the 6 minutes immediately following placement of the fish.

For the novel tank, test fish were placed in a tank 23 cm × 6 cm, 12 cm water depth, divided into three 4 cm zones) and their time spent in the different depth zones recorded for 6 minutes immediately after the fish were placed.

For the shoaling test, groups of four fish of either sex were placed into a large tank (45.5 cm ×25 cm, water depth 8 cm) and their swimming recorded for 6 min after 2 min of acclimatization time. Fish were simultaneously tracked and the pairwise Euclidean distance between each pair of fish determined and averaged per frame using commercially available Actual Track software (Actual Analytics, Edinburgh).

To test mating success, pairs of fish were placed into mating tanks (17.5 cm × 10 cm, water depth 6 cm) with a transparent divider in the evening. The next morning the divider was pulled at lights-on and the fish were allowed to breed for 1 hour. Each pair was bred 4 times every other day. Numbers of fertilized eggs in the clutch and the percentage of successful matings were recorded. A mating attempt was sored as successful, when fertilised eggs were produced.

### Statistical analyses

Quantitative data were tested for normality (Shapiro-Wilk test, *p< 0.05) and heteroscedasticity (Levene's test, *p < 0.05) to determine types of statistical comparisons. Variability of values is always given as SEM.Statistical significance was determined using Student's t-test for parametric data (with Welch's correction for heteroscedastic data) or Mann-Whitney U-test for nonparametric data. For multiple comparisons, we used one-way ANOVA with Bonferroni's post-hoc test for parametric homoscedastic data, one-way ANOVA with Welch's correction and Games-Howell post-hoc test for heteroscedastic data, and Kruskall-Wallis test with Dunn's post-test for nonparametric data. The shape of distributions was assessed using a Kolmogorov-Smirnov test (Fig.10). Randomisation was performed by alternating allocation of fish between control and treatment groups. No experimental animals were excluded from analysis. All relevant data are available from the authors.

## RESULTS

### Intraventricular injection of 6OHDA ablates specific populations of dopaminergic neurons and locally activates microglia.

To ablate dopaminergic and noradrenergic (Th^+^) neurons in the absence of damage to tissue and ERG processes, we established an ablation paradigm that relies on intraventricular injections of 6OHDA. Of the quantifiable Th^+^ cell populations in the brain ^33^, we found no effect of 6OHDA injection on cell numbers in populations 2, 7, 9, 10, 13 and 15/16 (data not shown). However, there was a 51% loss in population 5/6 (control: 484 ± 24 cell profiles; 6OHDA: 235 ± 14 cell profiles), 19% loss of TH^+^ cells in population 11 (288 ± 12 in controls vs. 234 ±16 in treated), 96% in population 12 (28 ± 1 in controls vs. 1 ± 0 in treated) and complete loss of noradrenergic neurons in the locus coeruleus (LC; 18 ± 1 in controls vs. zero in treated; Fig. 1A,B). Higher doses of 6OHDA did not increase loss of Th^+^ cells (data not shown). Consistent with Th^+^ cell loss, we found a 45% reduction in dopamine levels, but no effect on serotonin or its metabolites after 6OHDA injection in the whole brain by HPLC (Fig. 1E). There were no obvious correlations between the distance of neurons from the injection site or morphology of the neurons and rates of ablation (see Fig 1A). Hence, we devised an ablation paradigm in which neurons in populations 5/6, 11, 12 and the LC were selectively vulnerable to 6OHDA.

**Fig. 1.**
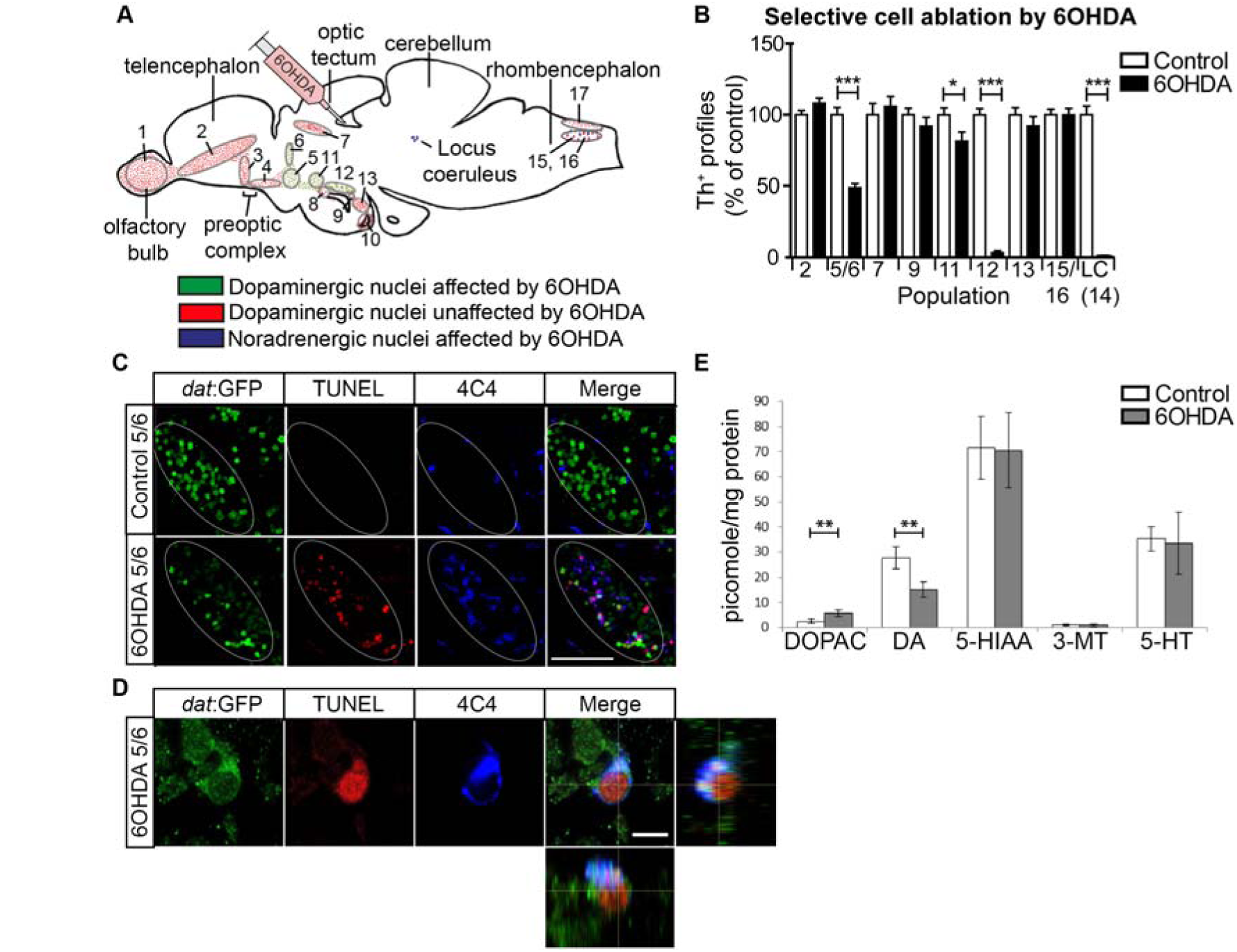
Specific populations of Th^+^ neurons are ablated by 6OHDA. **A**: A schematic sagittal section of the adult brain is shown with the 6OHDA resistant dopaminergic cell populations (red) and the vulnerable dopaminergic (green) and noradrenergic populations (purple) in relation to the injection site in the third ventricle indicated. **B**: Quantification of cell loss after toxin injection at 2 dpi is shown. **C**: Sagittal sections of population 5/6 are shown in a dat:GFP transgenic fish. This shows elevated TUNEL and microglia labelling in population 5/6 after ablation. Note that areas of elevated TUNEL and microglial labelling follow the outlines of the dat:GFP^+^ cell population (ellipse) in the 6OHDA treated animals, but not controls, indicating localised labelling.D: A high magnification is shown of a TUNEL^+^/dat:GFP^+^ dopaminergic neuron thatis engulfed by a 4C4^+^ microglial process (lateral and orthogonal views).**E**: Injection of the toxin decreases levels of dopamine (DA), increases levels of the metabolite DOPAC, but leaves serotonin (5-HT) and metabolites (5-HIAA, 3-MT) unaffected, as shown by HPLC. Student’s T-test (with Welch’s correction for heteroscedastic data) and Mann Whitney-U tests were used for pairwise comparisons in B and D (*p < 0.05; ** p < 0.01; 367151 p < 0.001). Bar in C = 50 μm, in D = 5 μm.

To determine whether 6OHDA injections led to specific death of Th^+^ neurons and activation of an immune response, we combined TUNEL labelling and immunohistochemistry for microglia using the 4C4 antibody, which selectively labels microglial cells ^34,37^ in a reporter fish for dopaminergic neurons *(dat:GFP)* ^28^ at 12 h post-injection. This indicated selective appearance of TUNEL^+^/*dat;GFP*+ profiles in the vulnerable populations, but not in the non-ablated populations or in areas not labelled by the transgene. Moreover, the density of microglial cells was selectively increased in these areas and some microglial cells engulfed TUNEL^+^/*dat;GFP*^+^ profiles, indicating activation of microglia (Fig. 1C,D; see also Fig. 8A for localised microglia reaction after 6OHDA treatment). Hence, 6OHDA only leads to death of circumscribed dopaminergic cell populations and elicits a localised microglial response.

**Fig. 8.**
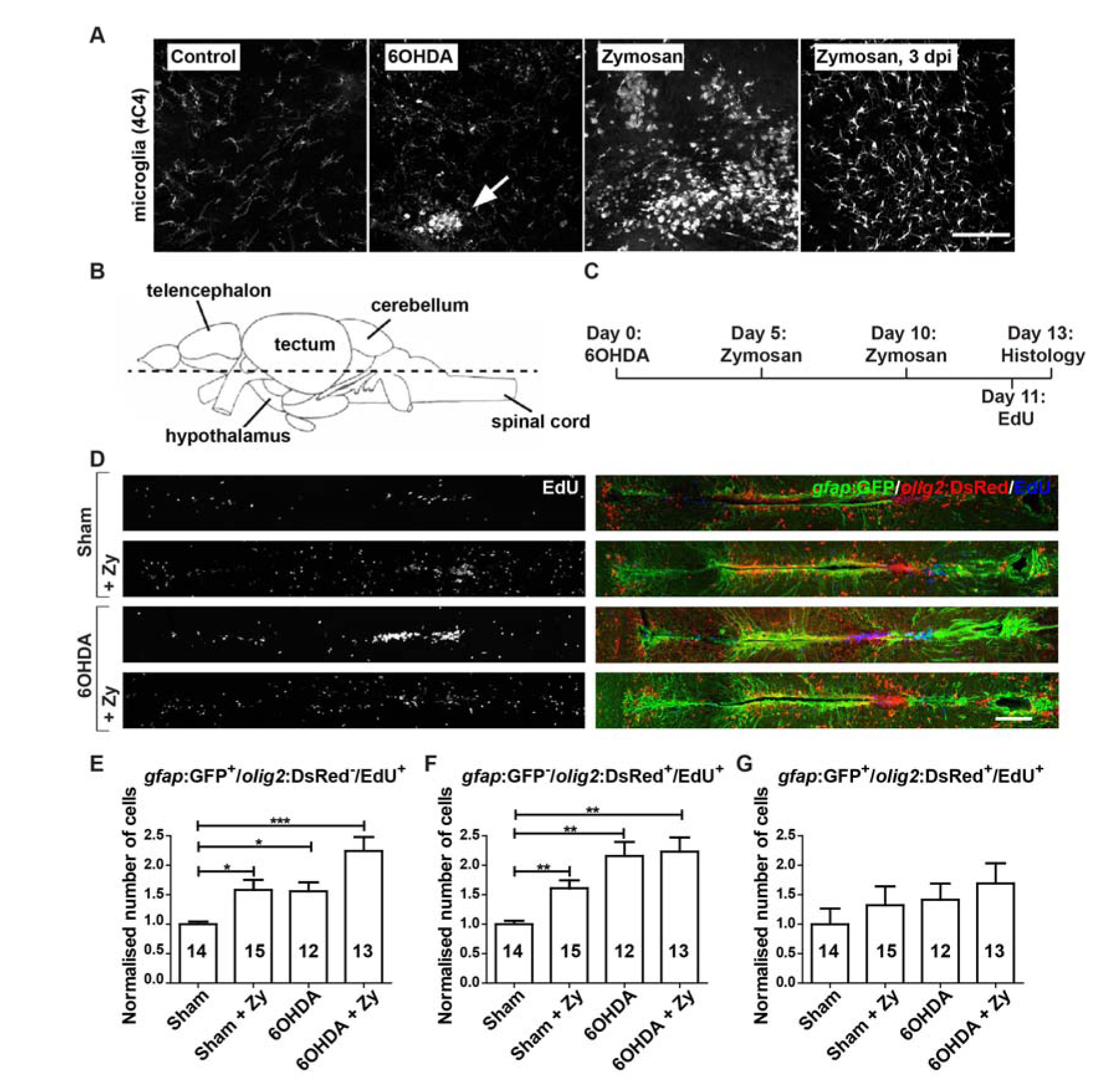
6OHDA and Zymosan injections both increase ERG proliferation. **A**: In sagittal sections of population 5/6 restricted microglia activation at 24 hours after 6OHDA injection (arrow in B) compared to sham-injected control animals (A) is observed. Zymosan injection leads to much stronger non-localised microglia response that lasts for at least three days (C,D). **B**: The horizontal plane of sectioning for G is shown; rostral is left for all panels. **C**: The experimental timeline for D-G is given. **D-G**: 6OHDA injections and Zymosan injections, alone or in combination with 6OHDA, increase proliferation of ERGs (D). This is true for only *gfap*:GFP^+^ (E) and only *olig2*:DsRed^+^ (F), but not double-labelled ERGs (G). One-way ANOVA with Welch’s correction and Games-Howell post-hoc test for H, I (*p < 0.05, **p < 0.01, ***p< 0.001), oneway ANOVA for J (p > 0.05). Scale bar in D = 100 μm for A-D; in G = 100 μm.

### Cell replacement and reinnervation patterns differ between dopaminergic cell populations.

To analyse whether lost Th^+^ neurons were replaced, we assessed Th^+^ cell numbers relative to controls without ablation for up to 540 days (1.5 years) post-injection of the toxin (dpi). The relatively small loss of cells in population 11 was compensated for at 42 dpi (not shown). In population 5/6, numbers were increased compared to 2 dpi, but were still lower than in controls by 42 dpi. However, at 180 dpi Th^+^ cell numbers were even slightly increased over controls. At 540 dpi numbers were similar to age-matched controls (Fig. 2A-E). In contrast, in population 12 and the LC, Th^+^ neuron numbers were never fully recovered. There was a small and transient recovery in cell numbers in these populations at 42 dpi, but by 540 dpi there were hardly any neurons present in population 12 and the LC (Fig. 2F-O). This indicates differential potential for cell replacement for different populations of dopaminergic neurons.

**Fig. 2.**
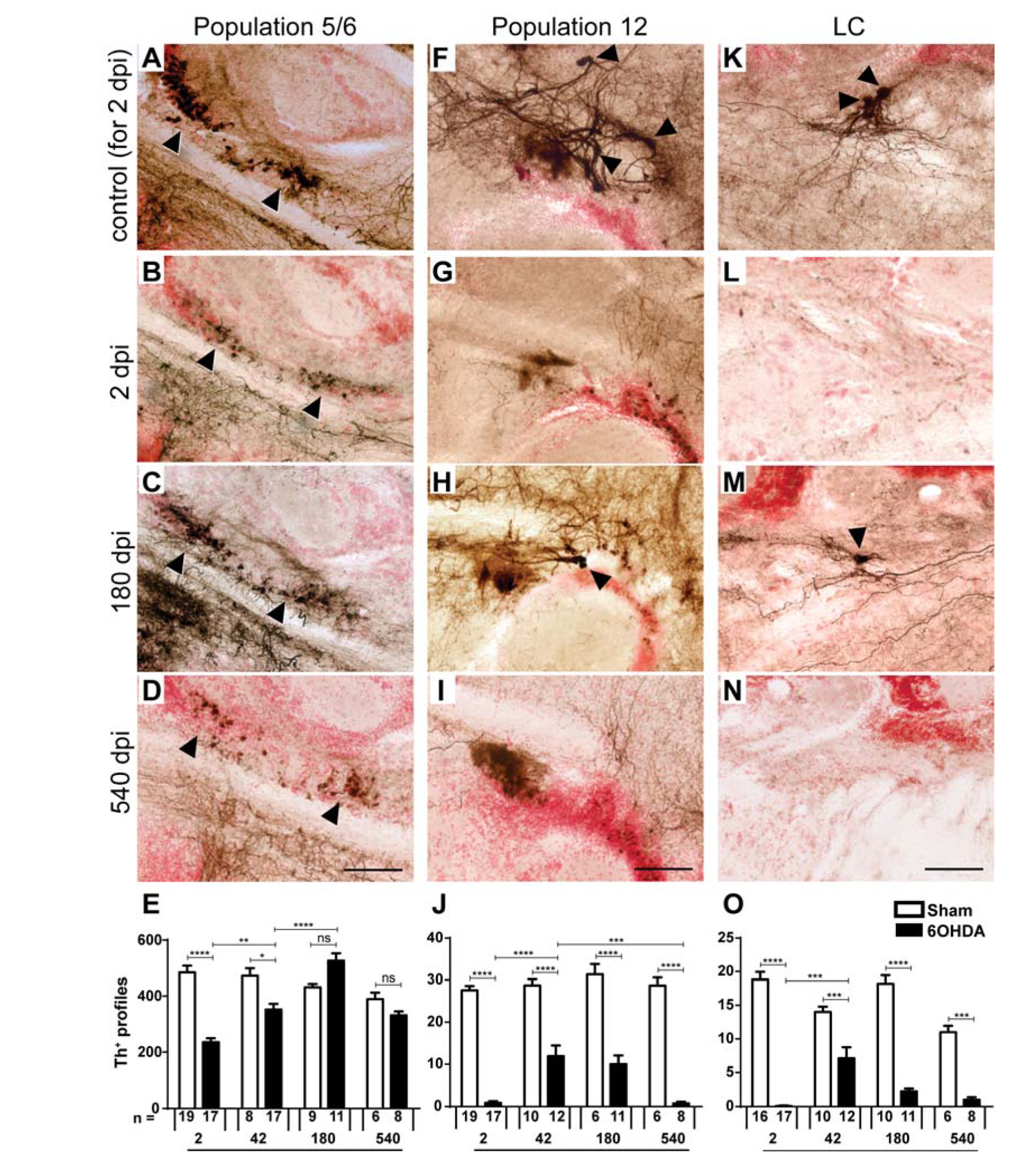
Replacement of Th^+^ neurons differs between brain nuclei. Sagittal brain sections are shown; dorsal is up, rostral is left. Some Th^+^ cell bodies are indicated by arrowheads. **A-E**: In population 5/6 the number of Th^+^ cells is reduced after toxin-induced ablation and back to levels seen in controls without ablation by 180 dpi. **F-J**: In population 12, a partial and transient recovery in the number of Th^+^ cells was observed at 42 dpl. **K-O**: In the LC there was also a partial and transient recovery of Th^+^ cell number. Note that example photomicrographs of controls are only shown for 2 dpi for clarity reasons, but all statistics were done with age-matched controls. Two-way ANOVA (p < 0.0001) with Bonferroni post-hoc test (*p < 0.05, **p < 0.01, 367151p < 0.001, 367151*p < 0.0001) for E, J, and O. Bars = 50 μm.

To determine whether restored dopaminergic neurons re-innervated their former target areas, we analysed a terminal field ventral to the predominantly locally projecting population 5/6 ^10^, which showed regeneration of cell bodies. After ablation, the density of Th^+^ innervation of this terminal field, measured semi-quantitatively by relative labelling intensity, was significantly reduced, compared to controls. This was still the case at 180 dpi, even though cell replacement had been almost completed by 42 days dpi. However, at 540 dpi, the axon density in 6OHDA-injected fish appeared not different from that in vehicle-injected controls anymore. This suggests slow restoration of local projections (Fig. 3A-E).

**Fig. 3.**
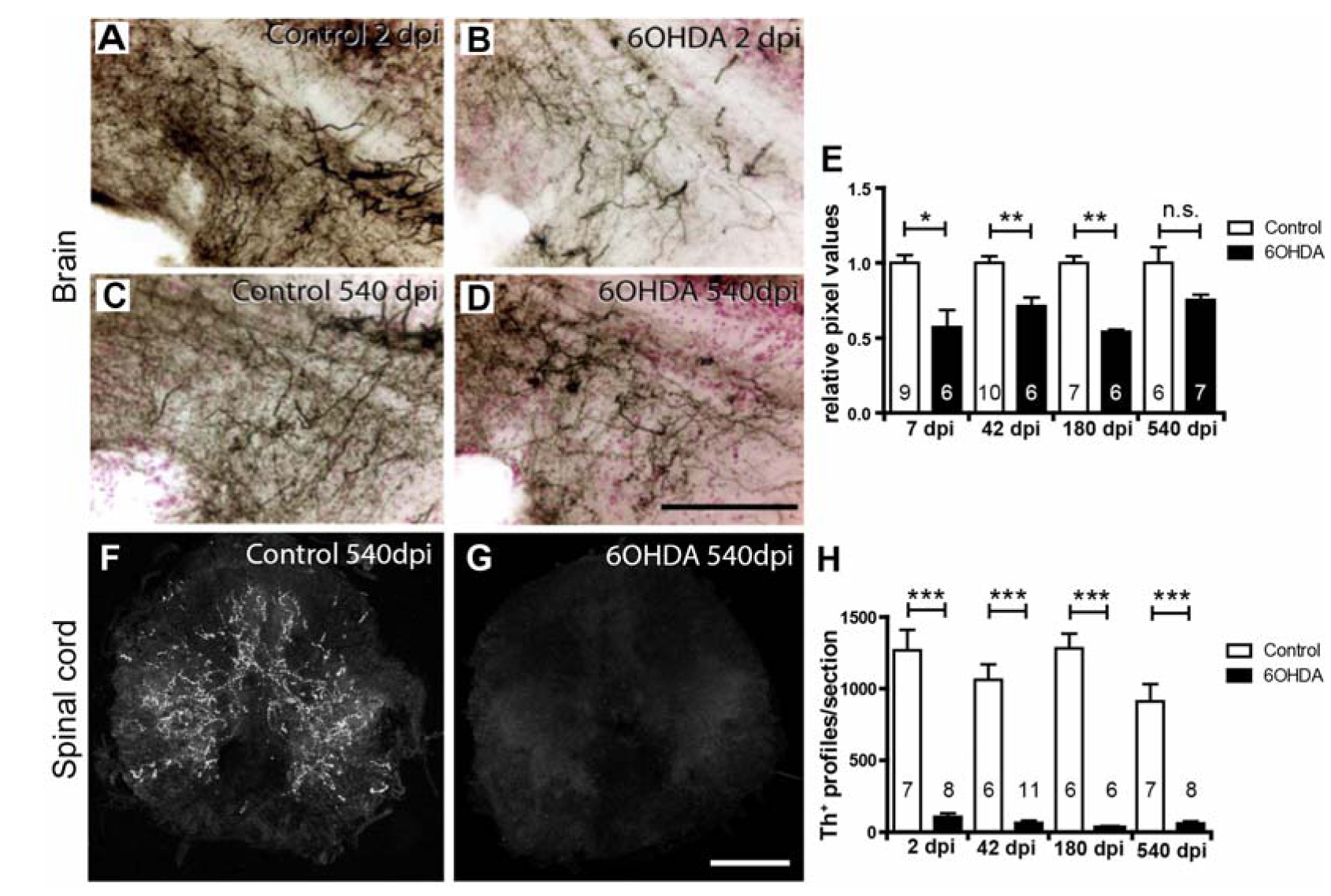
Th+ axons are inefficiently regenerated. **A-D**: Immunohistochemical detection of Th^+^ axons (black on red counterstain) in sagittal sections through a terminal field of TH^+^ axons ventral to population 5/6 is shown. Compared to controls **(A),** density of these axons is reduced at 2 dpi (**B**), and is more similar to age-matched controls (C,D) at 540 dpi. **E**: Semi-quantitative assessment of labelling intensity in the area depicted in D-G indicates significant loss of innervation at all time points except the latest, 540 dpi. **F,G**: Spinal cross sections are shown. Compared to age-matched controls (A, immunofluorescence for Th is very low at 540 dpi **(B). H**: Quantification of spinal Th^+^ axons indicates a lack of regeneration of the spinal projection. Student’s T-tests (with Welch’s correction for heteroscedastic data) or Mann-Whitney U tests were used for pairwise comparisons as appropriate (*p <0.05; ** 0.01; * * * p < 0.001). Bar in D = 100 μm for A-D; bar in G = 100 pm for F,G.

Since population 12 and the LC, which show little cell replacement, provide all Th^+^ innervation to the spinal cord ^12-14^, we assessed innervation of Th^+^ axons of the spinal cord. In animals without ablation, we always observed Th^+^ axons in the spinal cord at a midthoracic level (n = 26). Between 2 and 540 dpi, these axons were extremely rare in the spinal cord in 6OHDA injected animals (Fig. 3F-H). Hence the Th^+^ projection to the spinal cord was ablated by 6OHDA treatment and not regenerated.

### Capacity for enhanced addition of new dopaminergic neurons after ablation correlates with presence of constitutive neurogenesis for different populations

To determine how dopaminergic neurons were replaced after ablation, we assessed whether neurogenesis of dopaminergic neurons could be observed and whether ablation of dopaminergic neurons changed generation rates. To that aim, we injected EdU daily for 7 days after 6OHDA injection, to maximise progenitor labelling. We analysed the number of Th^+^/EdU^+^ neurons at 6 weeks post-injection, allowing sufficient time for differentiation of Th^+^ neurons (Fig. 4A). Even in the non-ablated situation, a low number of double-labelled neurons was observed in populations that were capable of neuron replacement, that is in populations 5/6, 8, and 11 (Fig. 4B,E-G). This indicates that dopaminergic neurons are constantly added to specific populations at a low rate.

After ablation, the number of double-labelled cells was increased 4.9fold in population 5/6 (Fig. 4C,D,E), compared to sham-injected animals. This was statistically significant. A similar non-significant trend was present in populations 8 and 11 (Fig. 4F,G). Hence, ablation of Th^+^ cells increases the rate of addition of new neurons to regenerating populations.

**Fig. 4.**
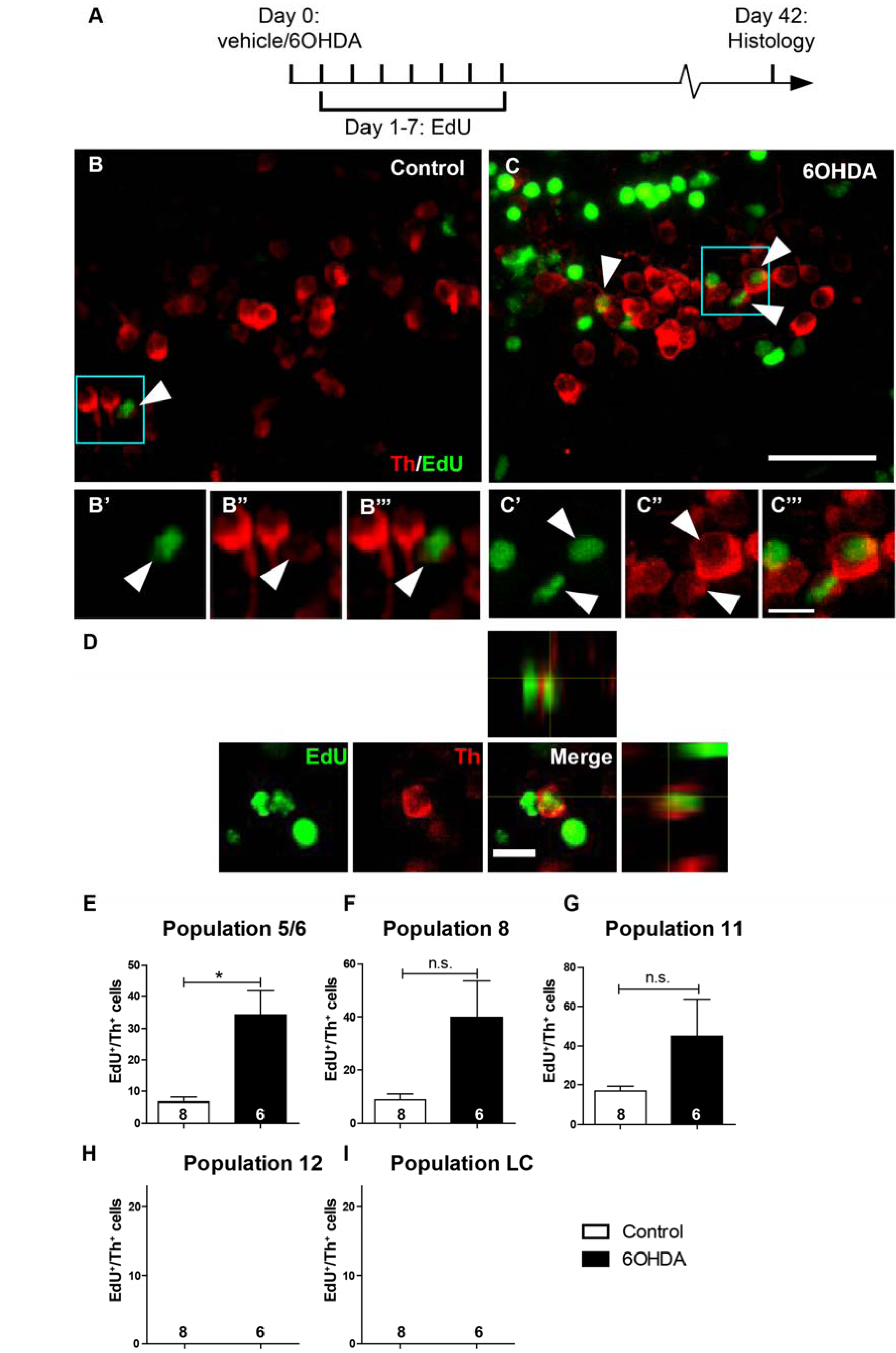
Generation of new Th^+^ cells is enhanced by prior ablation only in dopaminergic populations showing constitutive neurogenesis. **A**: The experimental timeline is given. **B,C**: In sagittal sections of population 5/6 (rostral left; dorsal up), EdU and Th double-labelled cells can be detected. Boxed areas are shown in higher magnifications in B’-C’’’, indicating cells with an EdU labelled nucleus, which is surrounded by a Th^+^ cytoplasm (arrowheads). **D**: A high magnification and orthogonal views of an EdU^+^/Th^+^ cell after 6OHDA treatment is shown. **E-I**: Quantifications indicate the presence of newly generated Th^+^ cells in specific dopaminergic brain nuclei (E-G). After 6OHDA treatment, a statistically significant increase in the number of these cells was observed for population 5/6. Note that population 12 and LC showed no constitutive or ablation-induced EdU labelled Th^+^ cells (H, I). (Student’s T-tests with Welch’s correction, *p <0.05,). Bar in C = 20 μm for A,B; bar in C ’’’ = 5μm for B’- C’’’, bar in D = 10 μm.

In contrast, in population 12 and the LC, which did not show strong replacement of Th^+^ neurons after 6OHDA injection in our histological analysis above, we did also not observe EdU^+^/Th^+^ neurons without or with ablation (Fig. 4H,I). Hence, differences in Th^+^ neuron replacement capacity correlate with differences in constitutive neurogenesis for distinct populations.

### New dopaminergic neurons are derived from ERGs

New Th^+^ cells are likely derived from local ERGs. The ventricle close to the 5/6 population is lined by cells with radial processes spanning the entire thickness of the brain. Most of these cells are labelled by *gfap:GFP*, indicating their ERG identity ^23^, and Th^+^ cells are located close to ERG processes (Fig.5A,B). Using PCNA labelling, we find that some of ERGs proliferate in the untreated brain, consistent with a function in maintaining dopaminergic and other cell populations (Fig. S3E,F).

**Fig. 5.**
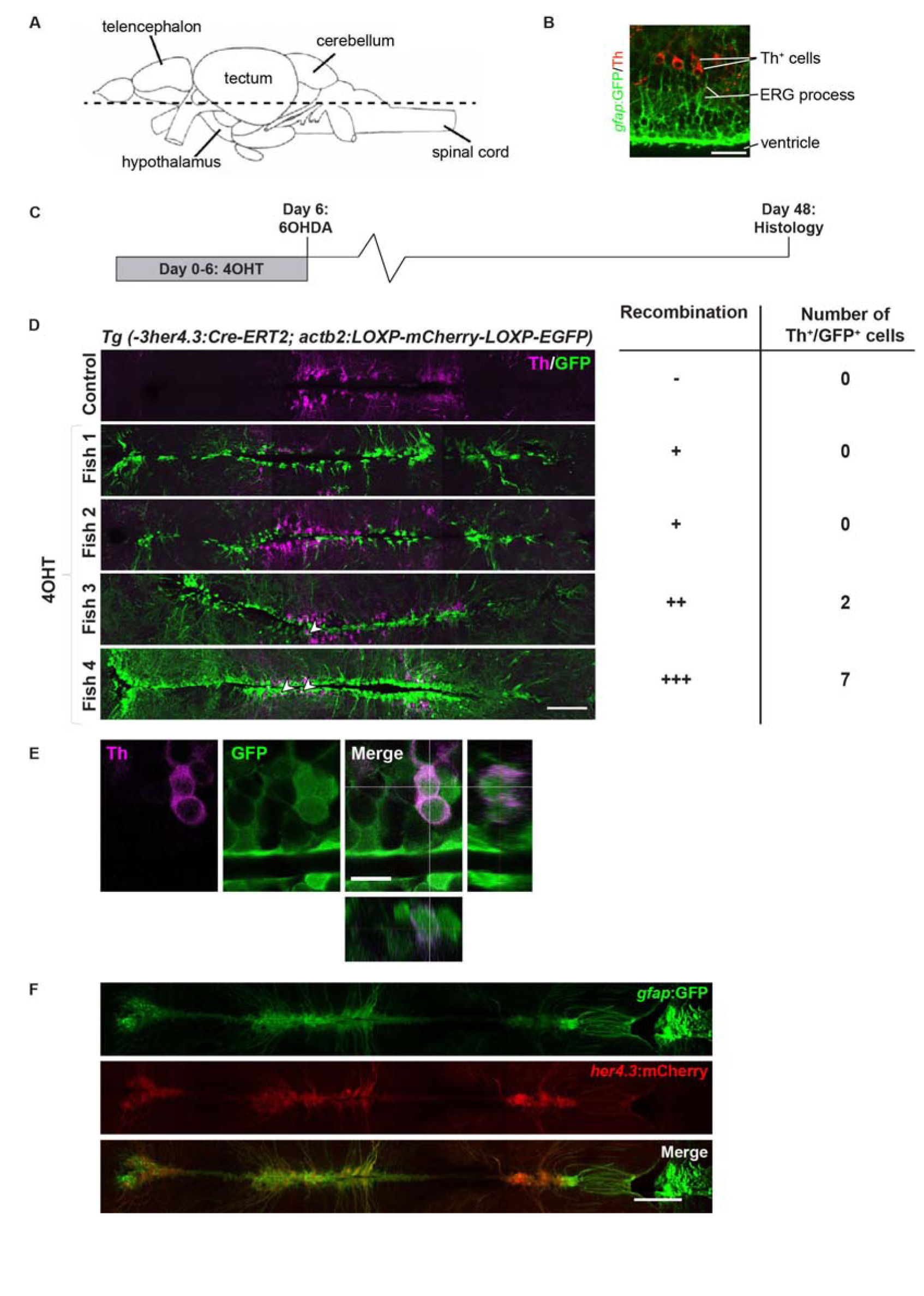
Genetic lineage tracing identifies ERGs as a source for new Th^+^ cells. **A**: The horizontal section level of all photomicrographs is indicated (rostral is left). **B**: Th^+^ cells are in close vicinity to *gfap*:GFP^+^ processes near the brain ventricle. **C,D**: Pulse-chase lineage tracing driven by *her4.3* indicates variable recombination and labelling of mainly ERGs and some Th^+^ neurons. **E**: High magnification and orthogonal views indicate Th^+^/GFP^+^ cells. **F**: her4.3:mCherry+ cells largely overlap with gfap:GFP labelling in double-transgenic animals. Scale bars: in B = 25 μm, D and F = 100 μm; in E = 10 μm.

To determine whether new Th^+^ cells are derived from ERGs, we used genetic lineage tracing with a *Tg(-3her4.3:Cre-ERT2)* x *Tg(actb2:LOXP-mCherry-LOXP-EGFP)* double-transgenic fish ^30^. In this fish, tamoxifen-inducible Cre is driven by the regulatory sequences of the *her4.3* gene. *her4.3* is specifically expressed in zebrafish ERGs ^38^. The second transgene leads to expression of GFP in ERGs and their progeny after Cre-recombination. We found a strong overlap between *gfap*:GFP and *her4.3*:mCherry labelling, indicating that the driver targets the appropriate cell population (Fig. 5F).

We incubated animals in tamoxifen for 6 days to induce recombination in ERGs, injected 6OHDA and waited for another 42 days for histological analysis. In animals without previous tamoxifen application, we did not observe any GFP^+^ cells. In tamoxifen-incubated animals, mostly ERGs were labelled at different densities, indicating variable recombination rates. In animals in which high recombination rates were achieved, we found GFP^+^/Th^+^ cells after the chase period, indicating that ERGs gave rise to dopaminergic neurons (Fig. 5C-E). However, we cannot exclude additional sources for new Th^+^ neurons that might be active during physiological or ablation-induced addition of these neurons.

### ERG proliferation is increased following ablation of dopaminergic neurons

To investigate whether ablation of dopaminergic neurons would lead to increased proliferation of ERGs, we determined EdU incorporation rates for different ERG populations (injected at 11 dpi and detected at 13 dpi; Fig. 6A,B). In the vicinity of the 5/6 population, most ERGs express *gfap.* Some of these co-express *olig2* and some express only *olig2,* as indicated by reporter fish double-transgenic for *gfap*:GFP and *olig2:DsRed* (Fig. 6D,E). ERGs that were only *gfap*:GFP^+^ showed increased rates of EdU incorporation after 6OHDA injection (Fig. 6F). Whereas ERGs that were only *olig2*:DsRed^+^ showed a similar trend (Fig. 6G), double-labelled ERGs did not show any 6OHDA-induced effect on proliferation (Fig. 6H). This indicates heterogeneity in the sensitivity of different ERG populations to dopaminergic cell ablation.

**Fig. 6.**
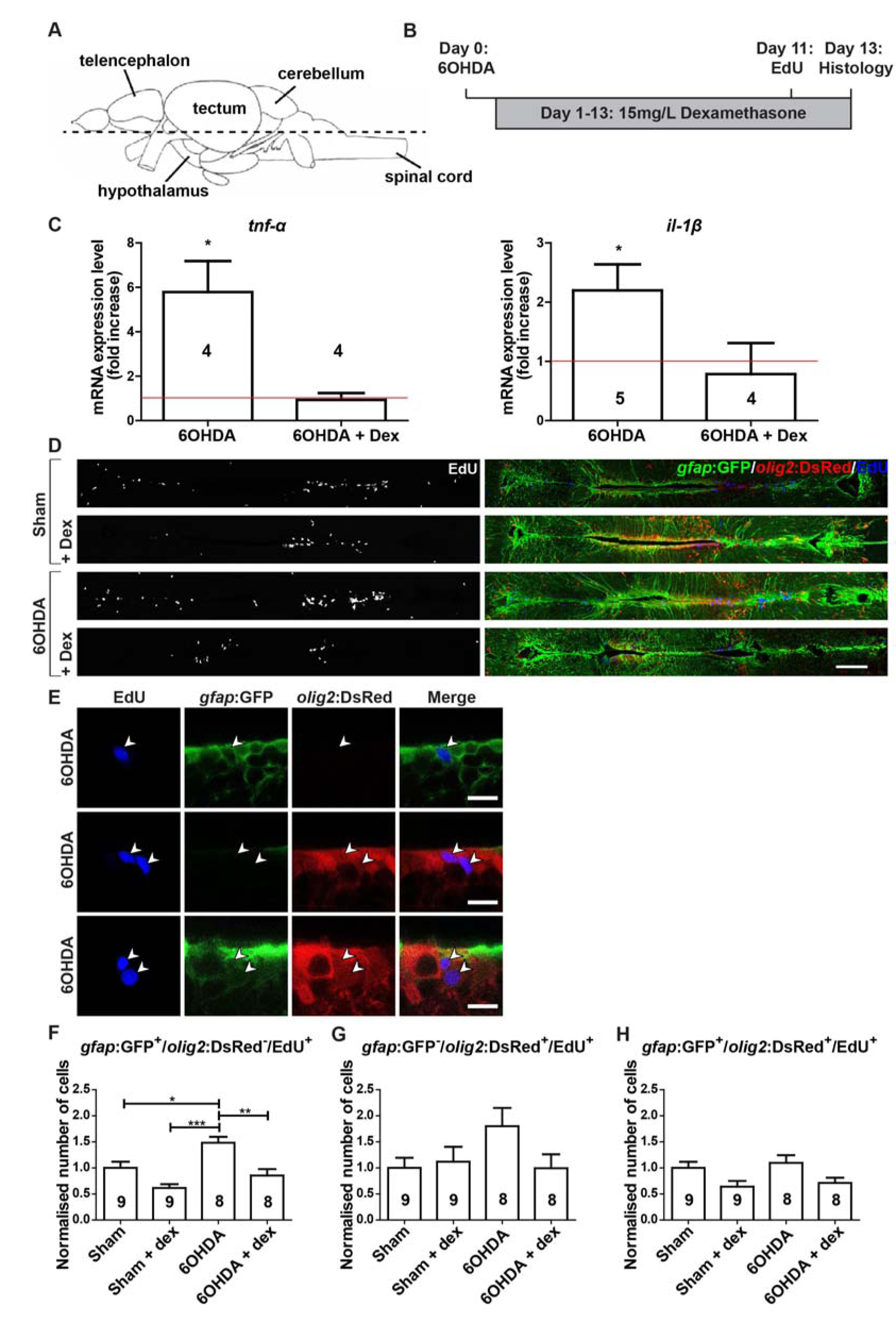
6OHDA injection increases ERG proliferation, which is abolished by dexamethasone treatment. **A,B**: The section level of photomicrographs (A, rostral is left) and experimental timeline (B) are given. **C**: Levels of *il-1β* and tnf-α are increased by 6OHDA treatment at 3 dpi, but not in the presence of dexamethasone, as shown by qRT-PCR. Each condition is normalised to sham-injected fish (shown by the red line; one-tailed one-sample t-tests, *p < 0.05). **D,E**: Overviews (D) of the quantification areas and higher magnifications of ventricular cells (E) are given. EdU-labels ERGs that are only gfap:GFP+, only *olig2:DsRed^+^* or both (arrowheads). **F-H**: The proliferation rate in only gfap:GFP^+^ ERGs is increased by 6OHDA injection and brought back to control levels by dexamethasone treatment (F). A similar non-significant trend is observed for only *olig2:GFP^+^* ERGs (G), but not for double-labelled ERGs (H). For F-H: One-way ANOVA with Bonferroni post-hoc test, *p < 0.05, **p < 0.01, *** p<0.001. Scale bars in C = 100 μm in D =10 μm.

To test whether reduced levels of dopamine after cell ablation (cf. Fig 1D) might trigger the increase in ERG proliferation, as in the salamander midbrain ^8^, we used extensive (see Material and Methods) injections of the dopamine D2-like receptor antagonist Haloperidol, which is effective in zebrafish ^18^, to mimic reduced dopamine levels in animals without ablation. However, this did not increase ventricular proliferation compared to sham-injected control animals (Fig. S2A-C), suggesting the possibility that reduced dopamine levels may not be sufficient to trigger progenitor cell proliferation. Taken together, the above observations support a scenario in which ablation of Th^+^ cells leads to enhanced generation of Th^+^ neurons mainly from *gfap*:GFP^+^ ERGs.

### Regeneration of Th^+^ cells depends on immune system activation

To test whether the observed activation of microglial cells (cf. Fig. 1C) was necessary for Th^+^ cell regeneration, we inhibited the immune reaction using dexamethasone bath application ^23^. qRT-PCR for principal pro-inflammatory cytokines *il-1beta* and *tnf-alpha* on horizontal brain sections comprising population 5/6, showed an ablation-induced increase in the expression of these cytokines in control fish that was consistent with the morphological activation of microglia. This increase was completely inhibited in the presence of dexamethasone, indicating that treatment was efficient (Fig. 6C).

Next, we determined if ERG proliferation was affected by dexamethasone incubation (for 14 days post-injection of 6OHDA, directly followed by analysis). Dexamethasone had no effect on proliferation rates of any ERG subtype in sham-injected controls, indicating that it did not influence ERG proliferation directly. In contrast, increased proliferation rates in only *gfap*:GFP^+^ ERGs of animals injected with 6OHDA were reduced to those seen in constitutive proliferation. This was statistically significant (Fig. 6F). ERGs that were only *olig2*:DsRed^+^ showed a similar trend (Fig. 6G). This showed that only ablation-induced proliferation of *gfap*:GFP^+^ ERGs depended on immune system activation.

To determine whether this early suppression of the immune response had consequences for the addition of newly generated Th^+^ cells to population 5/6, we incubated animals with dexamethasone for 14 days after ablation and analysed Th^+^ neuron addition at 42 days after ablation. This showed lower numbers of Th^+^/EdU^+^ neurons and lower overall numbers of Th^+^ neurons compared to 6OHDA treated animals without dexamethasone treatment (Fig. 7A-E).

**Fig. 7.**
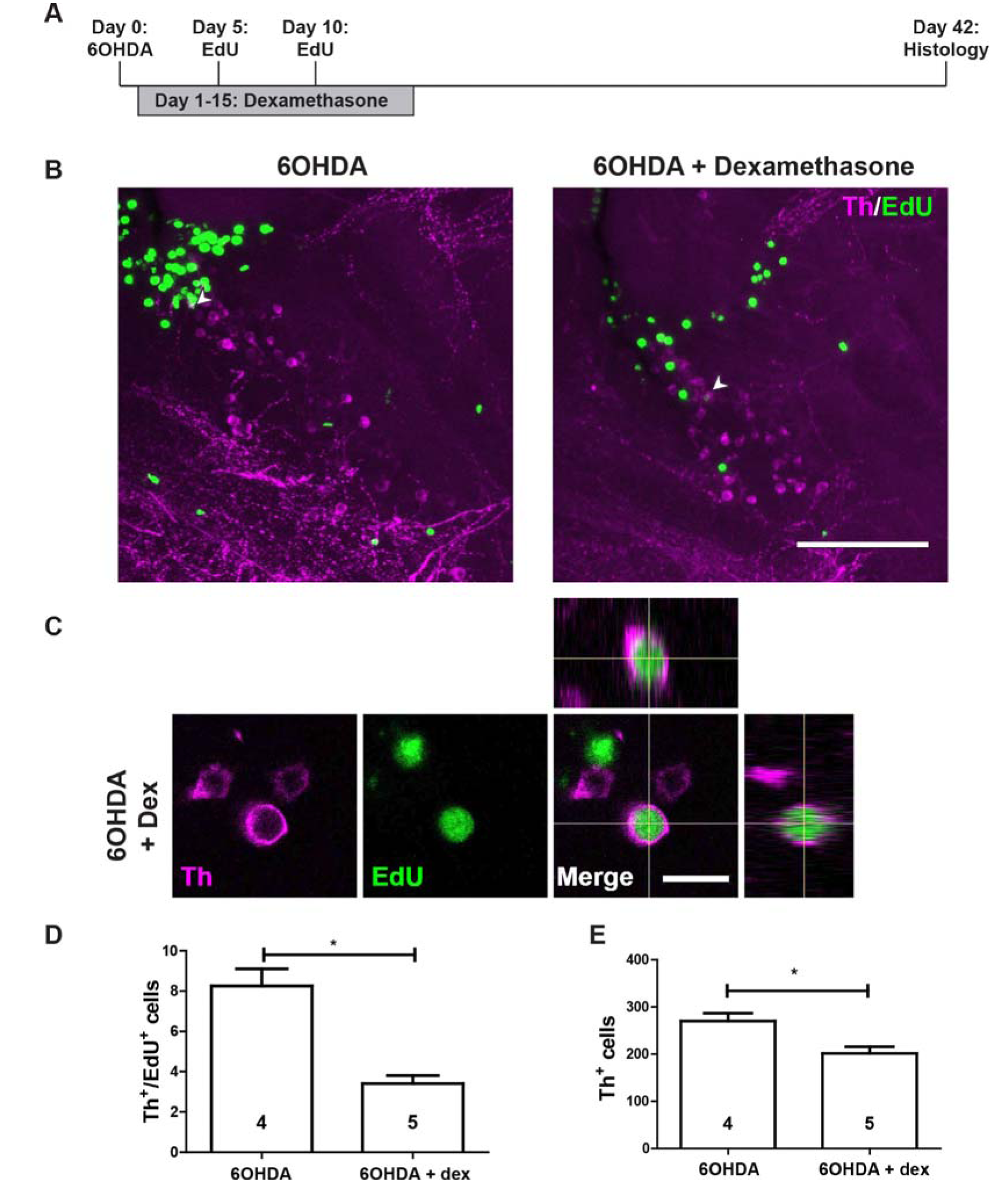
Dexamethasone inhibits regeneration of Th^+^ neurons in the 5/6 population. **A**: The experimental timeline is given. **B**: In sagittal sections of population 5/6, EdU^+^/Th^+^ neurons are present (arrowheads) after 6OHDA injection, with or without addition of dexamethasone. **C**: High magnification and orthogonal views of an EdU^+^/Th^+^ neuron are shown. **D,E**: The number of EdU^+^/Th^+^ (D; Mann Whitney-U test, *p < 0.05) and the overall number of Th^+^ neurons (E; Student’s t test, *p < 0.05) are reduced by treating 6OHDA-injected animals with dexamethasone. Scale bar in B = 100 pm; in C = 10 μm.

Next we asked whether dexamethasone treatment would reduce addition of new Th^+^ neurons that are constitutively added to the 5/6 population in the absence of ablation. Incubating fish with dexamethasone without 6OHDA injection did not alter the number of new Th^+^ neurons (Fig. S1A-E). The effect of dexamethasone on Th^+^ neuron addition only after 6OHDA treatment matched the effects of dexamethasone on ERG progenitor proliferation.

Hence, dexamethasone treatment early after ablation led to reduced rates of ERG proliferation and later Th^+^ neuron addition to population 5/6. This shows that most of regenerative neurogenesis depends on immune system activation.

### Augmenting the immune response enhances ERG proliferation, but not dopaminergic neuron regeneration

To determine whether the immune response was sufficient to induce dopaminergic cell generation and could be augmented to boost regeneration we used Zymosan A injections into the ventricle, compared to sham-injected controls and 6OHDA injection^23^. 6OHDA injection only led to local increase of 4C4 immunoreactivity, e.g. in the 5/6 population (Fig. 8A). In contrast, Zymosan injection led to a strong general increase in immunoreactivity for the microglia marker 4C4 that lasted for at least 3 days (Fig. 8A). Hence Zymosan injections can be used to boost the inflammatory reaction.

Without prior ablation of Th^+^ neurons, Zymosan injections led to increased proliferation of only *gfap:GFP*^+^ and only *olig2*:DsRed^+^ ERGs, but not of double-labelled ERGs, compared to untreated controls (Zymosan A injections at day 5 and 10 after 6OHDA injection, EdU application at 11 days post-injection, analysis at 13 days post-injection; Fig. 8B-G). After 6OHDA-mediated cell ablation, Zymosan treatment showed a trend to further enhance proliferation of only *gfap:GFP*^+^ ERGs compared to fish only treated with 6OHDA (Fig. 8E). However, this relatively weak additive effect was not statistically significant. Hence, Zymosan increased proliferation of mainly *gfap:GFP*^+^ ERGs independently of an ablation, and potentially slightly increased proliferation beyond levels induced by 6OHDA treatment alone.

To dissect whether the effect of immune system stimulation on ERG proliferation may have been mediated by the leukotriene LTC4, as in the mechanically injured telencephalon ^23^, we injected animals with the compound. This elicited a weak microglia response after 3 daily injections, as shown by 4C4 immunohistochemistry, but proliferation of ERGs was not altered (Fig. S3A,B). This suggests possible brain region-specific mechanisms of ERG proliferation.

To determine whether the increased ERG proliferation observed after Zymosan treatment alone would lead to generation of supernumerary Th^+^ neurons, we determined numbers of EdU^+^/Th^+^ and overall numbers of Th^+^ neurons at 42 days after a sham injection followed by two injections of Zymosan at 5 and 10 dpi. We did not observe any changes in these parameters (Fig. S1A-E), indicating that additional mechanisms may control dopaminergic differentiation of new cells.

To investigate whether Zymosan treatment was able to improve regeneration of Th^+^ neurons after ablation, we analysed the number of EdU^+^/Th^+^ and the total number of Th^+^ neurons after 6OHDA induced ablation, followed by Zymosan treatment, in the same experimental timeline as above. We did not observe any changes in EdU^+^/Th^+^ and overall numbers of Th+ cells compared to animals that only received 6OHDA injections (Fig. 9A-E). Hence, Zymosan treatment was sufficient to increase ERG proliferation but insufficient to boost regeneration of Th^+^ neurons.

**Fig. 9.**
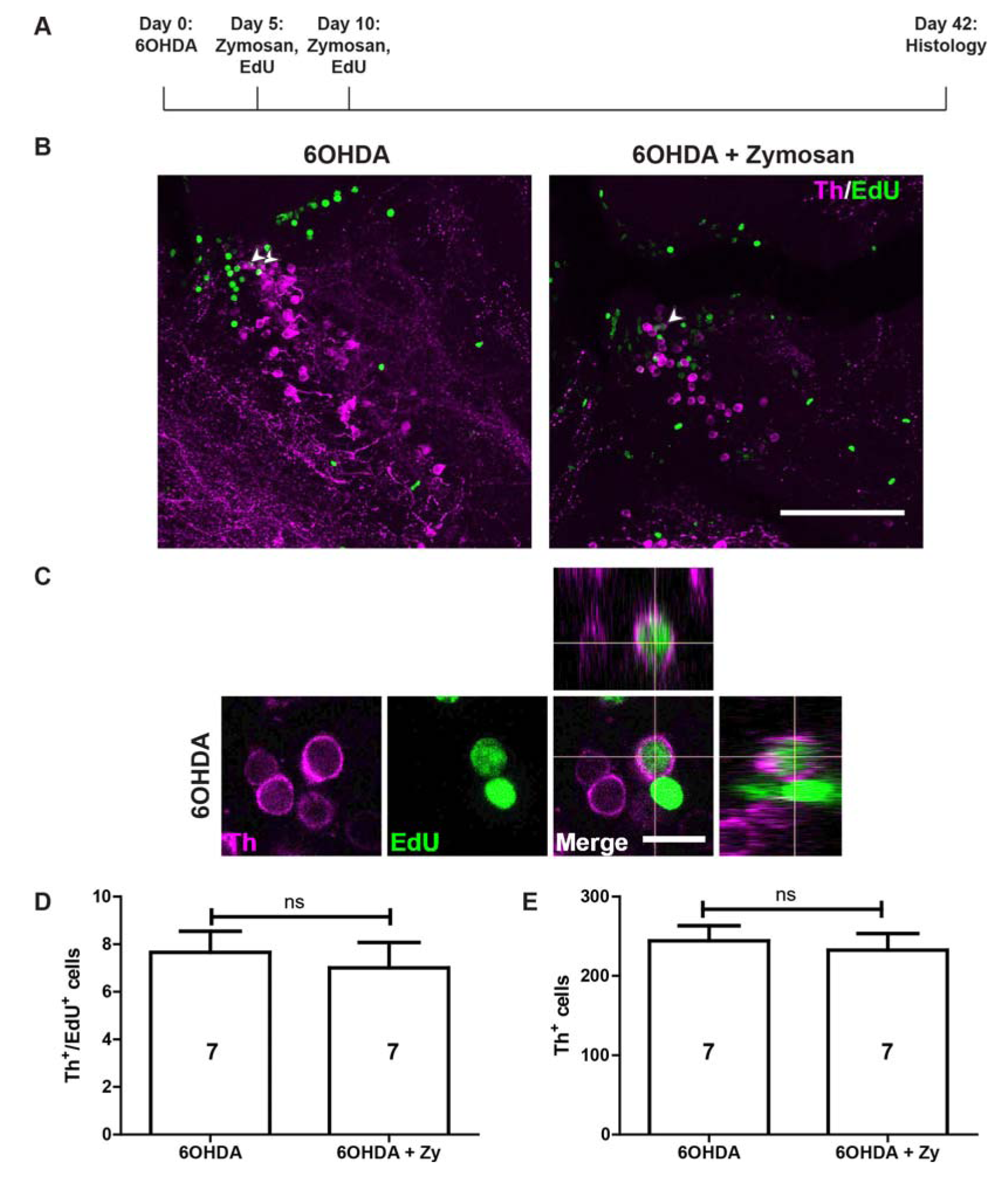
Zymosan treatment does not augment Th+ neuron replacement in population 5/6. **A**: The experimental timeline is indicated. **B**: In sagittal sections of population 5/6, EdU+/Th^+^ neurons can be observed (arrowheads) after 6OHDA injection with or without addition of Zymosan. **C**: High magnification and orthogonal views of an EdU^+^/Th^+^ neuron are shown**. D,E**:The number of EdU^+^/Th^+^ (D) and the overall number of Th^+^ neurons (E) are not increased by treating 6OHDA-injected animals with Zymosan (Student’s T-tests, p > 0.05). Scale bar in B = 100 μm; in C = 10 μm.

### Ablation of dopaminergic neurons leads to specific functional deficits

To determine whether loss of Th^+^ neurons had consequences for the behaviours of the fish, and whether these would be recovered after regeneration, we first recorded individual swimming activity in a round arena of fish that received 6OHDA injections and sham injections at 7 days after ablation. No differences were observed in the distance moved and velocity (average and frequency distribution) or the preference of fish for the periphery or inner zone of the arena (Fig. 10A-C and not shown) during the 6 minute observation period. This indicated that swimming capacity and patterns were not overtly affected by the ablation.

**Fig. 10:**
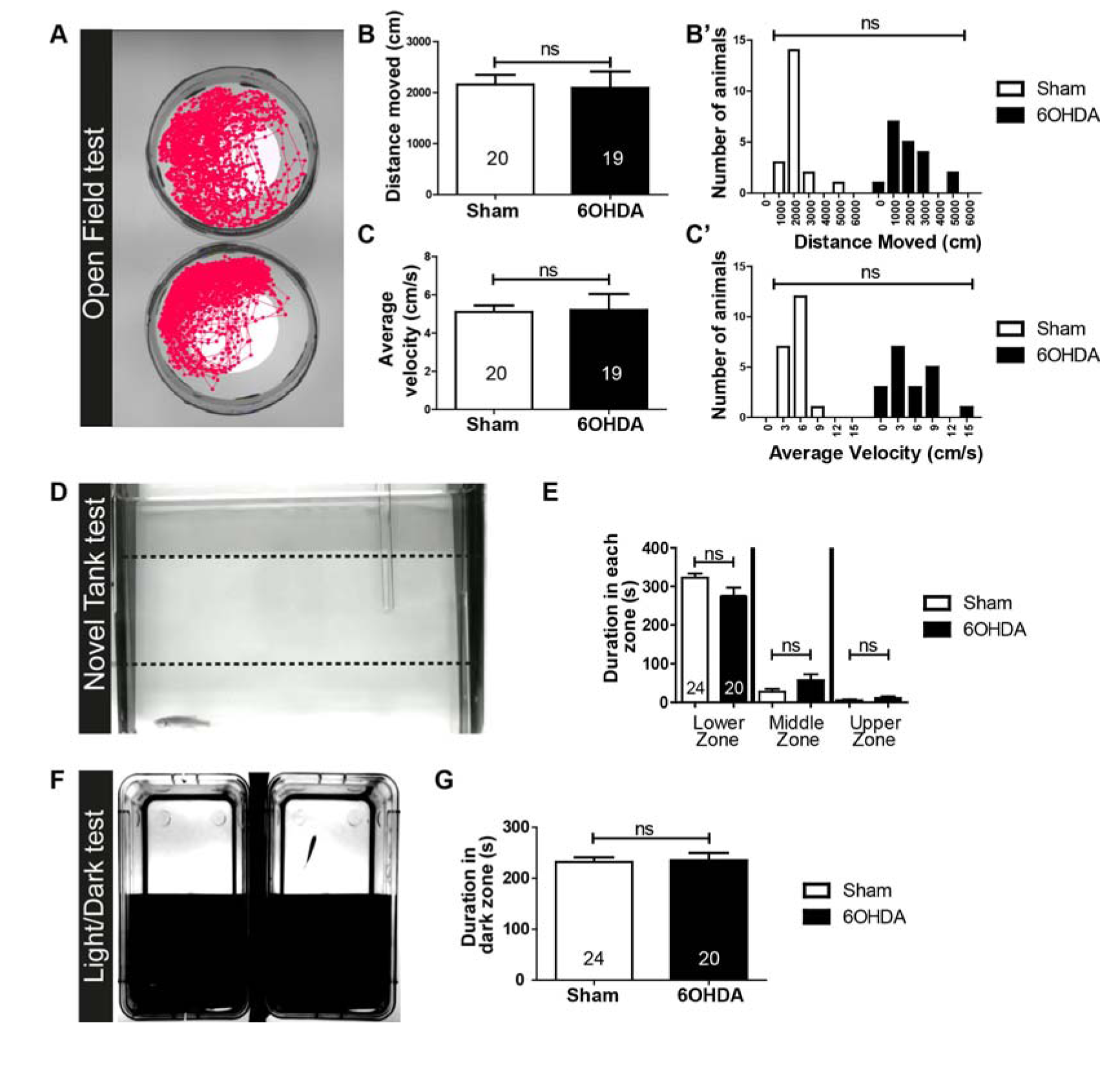
Basic swimming parameters and anxiety-like behaviours are unaltered by 6OHDA treatment.**A-C**: Typical swim tracks are shown (A). Quantifications of averages (B,C) and frequency distributions (B’,C’) of distance swum (B) and average velocity (C) showed no differences between control and treated fish (Student’s T-test, p > 0.05 for B and C; Kolmogorov-Smirnov Test, p > 0.05 for B’ and C’). **D,E**: A side view of a fish preferring the lower third of a novel tank is shown (D). Quantifications of time spent in the different depth of the tank (E) show the same preference for the lowest compartment in control and 6OHDA treated fish (Mann-Whitney U-tests, p > 0.05). **F**: The setup for the light/dark preference test is shown from above (F). Quantifications (G) indicate that control and treated fish do not differ in their preference for the dark compartment in a 300 seconds observation period (Student’s T-test, p > 0.05).

We used tests of anxiety-like behaviours, namely the novel tank test, in which fish initially prefer to stay at the bottom of the unfamiliar new tank, and the light/dark choice test ^39-41^, in which fish stay most of the time in the dark compartment. Indeed, fish in all groups showed strong preferences for the bottom of the tank or the dark compartment, respectively, indicating the expected behaviours. However, fish did not show any differences in behaviour after 6OHDA induced ablation of Th^+^ neurons (Fig. 10D-G). Hence, we could not detect effects of Th^+^ cell ablation on anxiety-like behaviours.

To test movement coordination, we analysed shoaling behaviour of the fish. Putting 4 fish together into a tank lets them exhibit shoaling, a natural behaviour to swim close to their conspecifics ^42^. This behaviour requires complex sensory-motor integration to keep the same average distance from each other. We found that shoals made up of fish treated with 6OHDA swam at an average inter-individual distance that was twice as large as that in control shoals at 7, 42 and 180 days post-injection (Fig. 11A,B). Hence ablation of Th^+^ neurons impaired shoaling behaviour and this behaviour was not recovered within 180 days dpi.

**Fig. 11.**
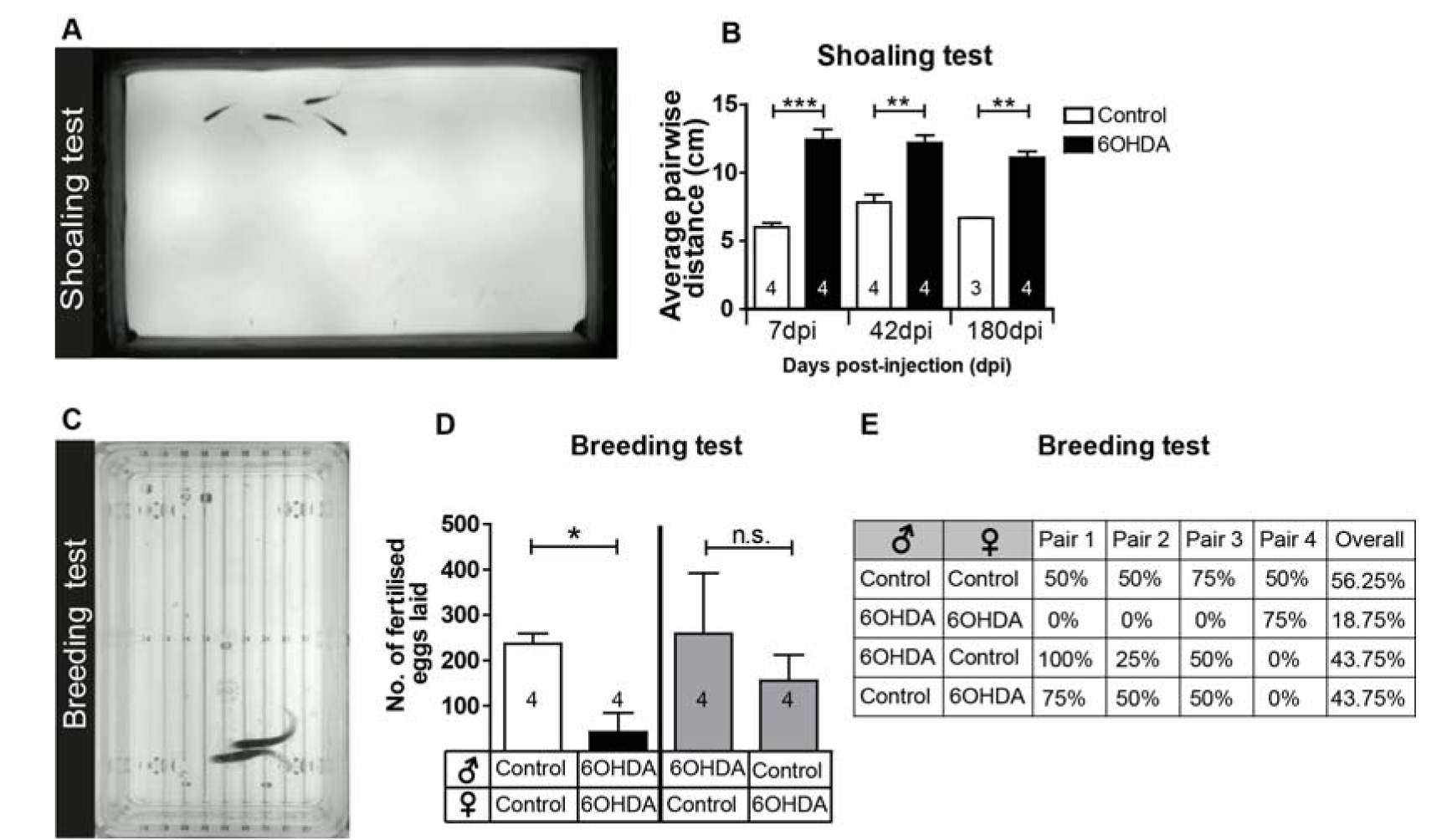
Injection of 6OHDA permanently impairs shoaling behaviour and decreases mating success. **A,B**: Shoaling behaviour as viewed from above in shallow water is shown in A. The average pairwise distance of fish from each other is significantly increased at all time points tested (Student’s T-tests with Welch’s correction for heteroscedastic data were used for pairwise comparisons; *p < 0,05, **p < 0.01, ***p < 0.001). N-numbers indicate number of shoals of 4 fish each. **C-E**: Fish showing mating behaviour are shown from above (C). Clutch sizes (D) and mating success rates (E) are strongly reduced when animals are mated after 6OHDA injection. Mating the same females or males with the same control fish showed that mating success does not depend on lack of 6OHDA in either males or females alone. N-numbers indicate different pairs of fish (D; Mann-Whitney U-tests, P < 0.05).

We reasoned that if manoeuvring of fish was impaired by ablation of specific Th^+^ cells, mating behaviour, which requires coordinated swimming of a male and female, might also be affected. Alternatively, reproductive functions could directly be influenced by dopamine ^43^. Indeed, ablation of Th^+^ cells in both male and females led to a reduced rate of successful matings and 84% fewer fertilised eggs laid than in control pairs over four mating events. Combining the same control females with the 6OHDA treated males and vice versa allowed intermediate egg production and mating success in both groups, indicating that male or female reproductive functions were not selectively affected (Fig. 11C-E). Hence, mating success was only strongly impaired when both males and females lacked specific Th^+^ neurons. This supports the notion that swimming coordination was permanently affected by the lack of regeneration in population 12 and the LC.

## DISCUSSION

Our results show that after ablation, Th^+^ neurons in some populations are replaced by newly formed neurons. Th^+^ neurons are derived from specific ERGs, which increase proliferation after ablation in the adult zebrafish brain. This regeneration depends on immune system activation. In contrast, Th^+^ neuron populations with long spinal projections only show sparse and transient replacement of neurons and never recover their spinal projections. Consequently, deficits in shoaling and mating behaviours associated with these anatomical defects never recover (schematically summarized in Fig.12).

**Fig. 12.**
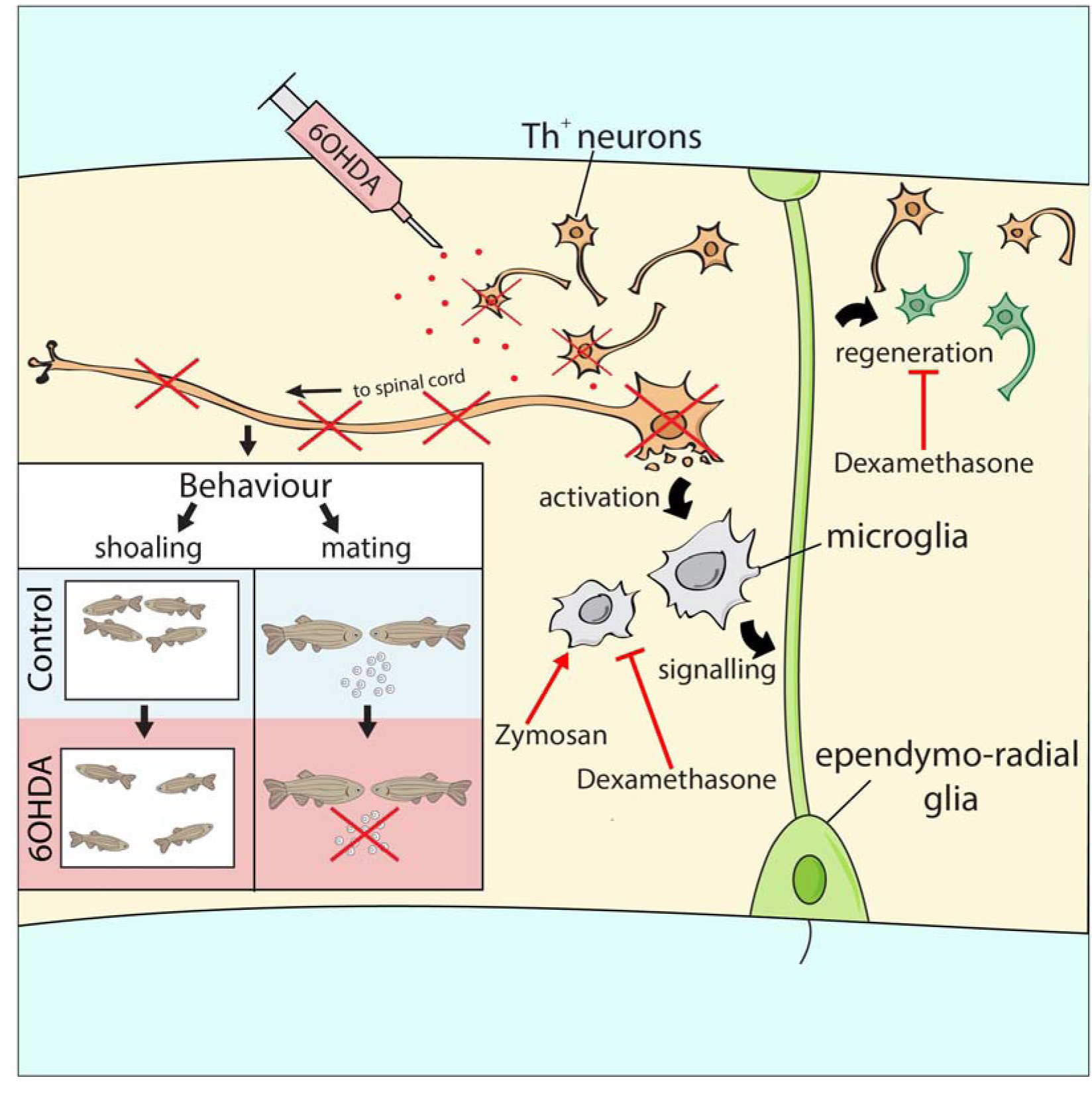
Schematic overview of results. 6OHDA injection ablates specific Th^+^ cell populations, leading to a microglia response, which is necessary for regeneration of new dopaminergic neurons from ERGs. This is blocked by dexamethasone, whereas Zymosan stimulates ERG proliferation, but not addition of Th^+^ neurons. Neurons projecting to the spinal cord are not replaced, associated with deficits in shoaling and mating behaviours.

### Th^+^neurons are regenerated from specific ERG progenitors after ablation

We observed a regenerative response after ablation of a subset of Th^+^ neurons, defined by an increased number of Th^+^ cells and ERGs labelled with a proliferation marker. Genetic lineage tracing showed that ERGs gave rise to at least some new Th^+^ neurons. However, we cannot exclude contributions from unknown progenitors or trans-differentiation of other neurons as a source for new dopaminergic neurons. Hence, ablation ofTh^+^ neurons is sufficient to elicit a regenerative reaction in ERG progenitor cells and protracted replacement of Th^+^ neurons.

Not all diencephalic ERGs may take part in regenerative neurogenesis of dopaminergic neurons. We find previously unreported heterogeneity in gene expression and proliferative behaviour. While *gfap*:GFP^+^ (overlapping with the ERGs labelled by genetic lineage tracing) and *olig2*:DsRed^+^ ERGs showed changes in proliferation in response to ablation or immune signal manipulation, those that expressed both transgenes were not altered in their proliferation rates by any of these manipulations, indicating that only specific ERGs may act as progenitor cells in a regeneration context.

In previous ablation experiments in larvae, different observations were made depending on the ablated cell populations. Either enhanced proliferation and replacement of neurons ^44^ or no reaction and long-term reduction in neuron number ^45^ has been reported. This underscores our findings that different populations of dopaminergic neurons are not regenerated to the same extent, even in larvae that show higher general proliferative activity than adults. Our observation supports that loss of Th^+^ cells leads to increased proliferation of progenitor cells and replacement of specific dopaminergic neuron populations.

### The immune response is necessary for regeneration of Th^+^ cells

We find that inhibiting the immune response after ablation leads to reduced proliferation in the ventricular zone and fewer new Th^+^ neurons. Interestingly, only ablation-induced ERG proliferation was affected by this treatment, consistent with findings for the zebrafish telencephalon ^23^. It has been proposed that different molecular mechanisms are involved in constitutive and regenerative neurogenesis ^46^. However, the immune-mediator LTC4, reported to promote the immune-dependent progenitor proliferation in the zebrafish telencephalon ^23^, did not elicit proliferation of ERGs in our experiments in the diencephalon, suggesting regional differences of immune to ERG signalling.

Alternatively, ERGs could be de-repressed in their activity by the observed reduction of dopamine levels in the brain. This has been demonstrated to be the case in the midbrain of salamanders ^8^. However, injecting haloperidol into untreated fish to mimic reduced levels of dopamine after ablation did not lead to increased ERG proliferation in the brain of zebrafish. This points to potential species-specific differences in the control of progenitor cell proliferation between zebrafish and salamanders.

Remarkably boosting the immune reaction with Zymosan was sufficient to enhance ERG proliferation, but was insufficient to increase number of new Th^+^ neurons in animals with and without prior ablation of Th^+^ neurons. This suggests that additional factors, not derived from the immune system, may be necessary for Th^+^ neuron differentiation and replacement.

### What are the reasons for differentia/ regeneration of dopaminergic neuron populations?

Constitutive neurogenesis we observe in specific brain nuclei correlates with regenerative success. For example, there is ongoing addition of Th^+^ cells in the regeneration-competent 5/6 population without any ablation, but this is not detectable in the non-regenerated populations 12 and LC. We speculate that in brain nuclei that constitutively integrate new neurons, factors that support integration of new neurons, such as neurotrophic factors and axon guidance molecules might be present, whereas these could have been developmentally down-regulated in populations that do not add new neurons in adults. Integration promoting factors may be rate-limiting for regeneration.

Alternatively, new neurons may fail to integrate into the network and perish. This may be pronounced for population 12 and the LC, which show complex axon projections ^10^. Some dopaminergic cells managed to repopulate population 12 and LC, but they did not persist. These populations have neurons with particularly long axons that are led by complex guidance molecule patterns, e.g. to the spinal cord during development ^47^. These patterns may have disappeared in adults and thus explain failure of these neurons to re-innervate the spinal cord. Some long-range axons can successfully navigate the adult zebrafish brain, such as regenerating optic axons ^48^, but particular populations of axons descending to the spinal cord do not readily regenerate ^49,50^ This correlates with constitutive neurogenesis in the optic system, but not in the descending brainstem projection.

### Specific ablation of circumscribed Th^+^ populations offers clues to their function

The long-lasting loss of about 28 dopaminergic neurons in population 12 and of 18 noradrenergic neurons in the LC is associated with highly specific functional deficits in shoaling and mating, but not overall locomotion or anxiety-like behaviours. Previous studies showed reduced overall locomotion after application of 6OHDA in adult zebrafish. However, in these studies, application routes were different, creating larger ablation in the brain ^7^ or peripheral rather than central lesions ^51^.

Among the lost neurons, population 12 contains the neurons that give rise to the evolutionarily conserved diencephalo-spinal tract, providing the entire dopaminergic innervation of the spinal cord in most vertebrates ^10^. Loss of this tract in larval zebrafish leads to hypo-locomotion, due to a reduction in the number of swimming bouts ^16,17^. Large scale ablation of diencephalic dopaminergic neurons in larvae also led to motor impairments ^52^. We speculate that in adults, dopamine in the spinal cord, which is almost completely missing after ablation, may modulate initiation of movement changes necessary for efficient shoaling and mating behaviour. However, descending dopaminergic projections also innervate the sensory lateral line ^17,53^. Altered sensation of water movements could thus also contribute to impaired ability to manoeuvre. Moreover, population 12 neurons have ascending projections ^10^ that could also be functionally important. We can also not exclude that some ablated dopaminergic neurons escaped our analysis but contributed to functional deficits.

Altered shoaling behaviour ^54^ and anxiety-like behaviour ^19,20^ has previously been correlated with alterations of the dopaminergic system, but not pinpointed to specific neuronal populations. Our results support that the fewer than 50 neurons that form the descending dopaminergic and noradrenergic projections are involved in shoaling behaviour, but not anxiety-like behaviour, as has been found for global manipulations of dopamine ^55^.

Dopamine-dependent behaviours can be recovered following regeneration of dopaminergic neurons. For example, in larval zebrafish, swimming frequency is normalised again after ablation and regeneration of hypothalamic dopaminergic neurons ^44^. In salamanders, amphetamine-inducible locomotion is recovered, correlated with regeneration of Th^+^ neurons after 6OHDA-mediated ablation ^56^. Here we show that regeneration of specific Th^+^ neurons that project to the spinal cord is surprisingly limited in adult zebrafish and not functionally compensated, which leads to permanent functional deficits in a generally regeneration-competent vertebrate.

## Conclusion

Specific Th^+^ neuronal populations in adult zebrafish show an unexpected heterogeneity in their capacity to be regenerated from specific progenitor populations. This system is useful to dissect mechanisms of successful and unsuccessful functional neuronal regeneration in the same model, and we show here that the immune response is critical for regeneration. Ultimately, manipulations of immune mechanisms in conjunction with pro-differentiation factors may be used to activate pro-regenerative mechanisms also in mammals to lead to generation and functional integration of new dopaminergic neurons.

## Acknowledgements

ACKNOWLEDGEMENTS

We thank Drs Bruce Appel, Marc Ekker, Daniel Goldman, and Pamela Raymond for transgenic fish, Joe Finney for data analysis and Stephen West for discussions. Supported by BBSRC (BB/M003892/1 to CGB and TB), an MRC DTG PhD studentship (to NOD), a BBSRC Eastbio PhD studentship (to LJC), and a grant from Sigrid Juselius Foundation to SS and PP.

## AUTHOR CONTRIBUTION

Conceptualization, NOD, LJC, CGB, and TB; Investigation, NOD, LJC, LC, SAS, KSM, PP and JDA; Writing: CGB, and TB.

---

**S1:**
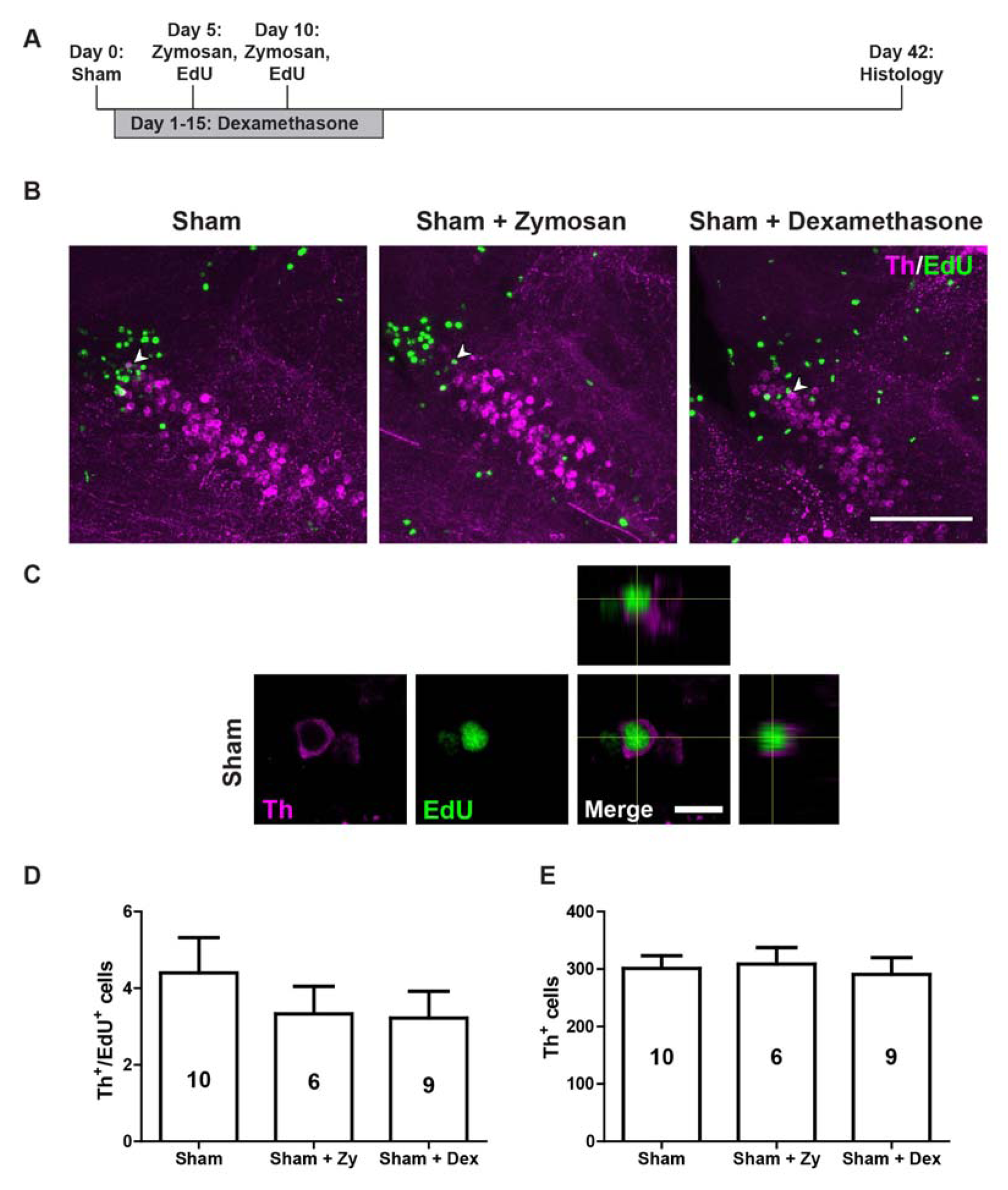
Immune system manipulations in animals in the absence of ablation do not influence addition of new Th^+^ cells. **A**: Timeline of experiments with either Zymosan or dexamethasone treatment. **B**: In sagittal sections of population 5/6, EdU^+^/Th^+^ neurons (arrowheads) can be observed in all experimental conditions. **C**: A high magnification and orthogonal views of a double-labelled neuron in a sham-injected animal are shown**. D,E**: In animals without ablation no changes are observed in the number of newly generated Th^+^ neurons and the overall number of Th neurons after dexamethasone or Zymosan treatment (One-way ANOVA with Bonferroni post-hoc test used in D and E, p > 0.05). Scale bar in B = 100 μm; in C = 10 μm.

**S2.**
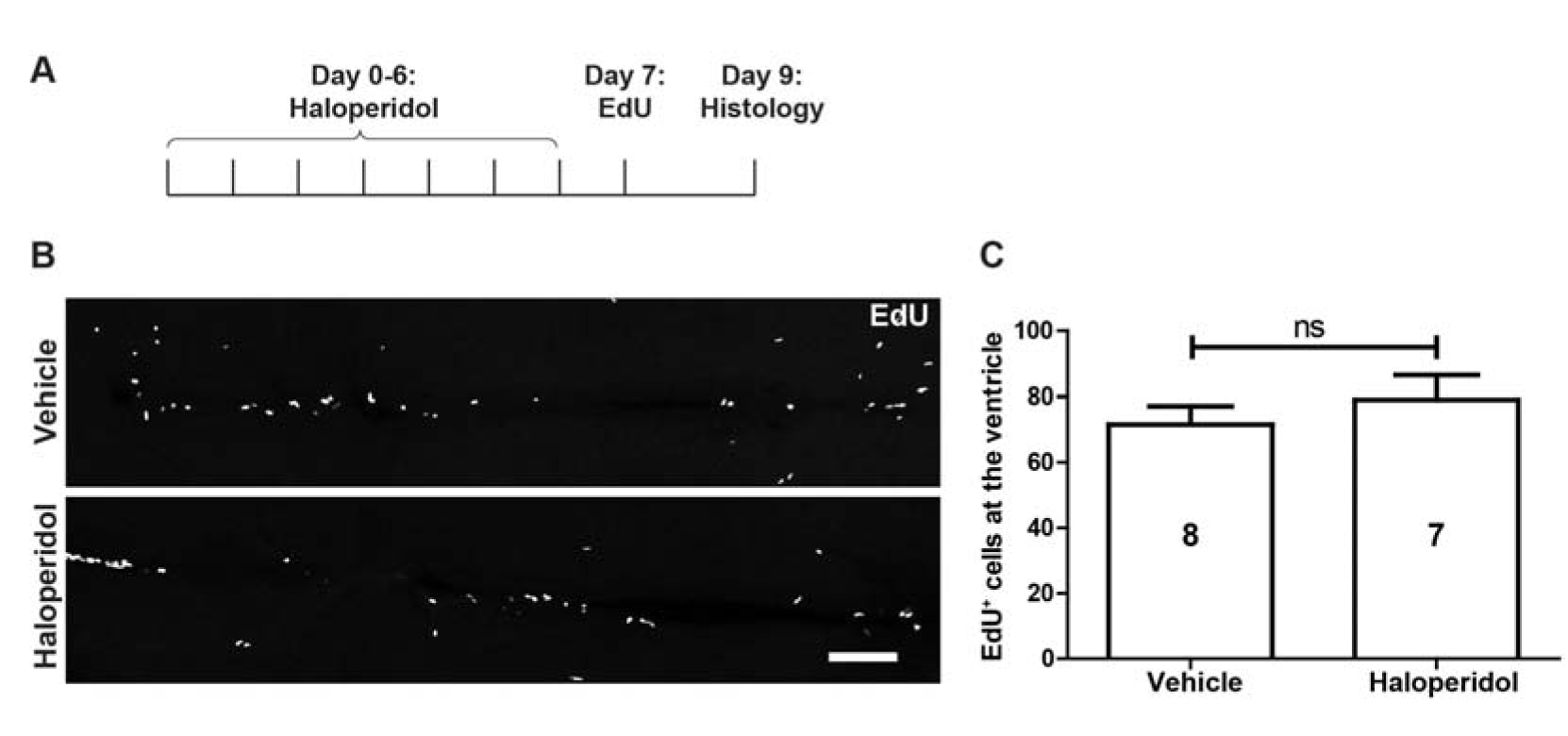
Inhibition of dopamine signalling does not affect ERG proliferation. **A**: The experimental timeline for B,C is shown. **B,C**: Horizontal sections (B) and quantification (C) show no effect of Haloperidol on proliferation in the ERG layer. (Student’s t-test, p > 0.05). Scale bar in B = 100 μm.

**S3:**
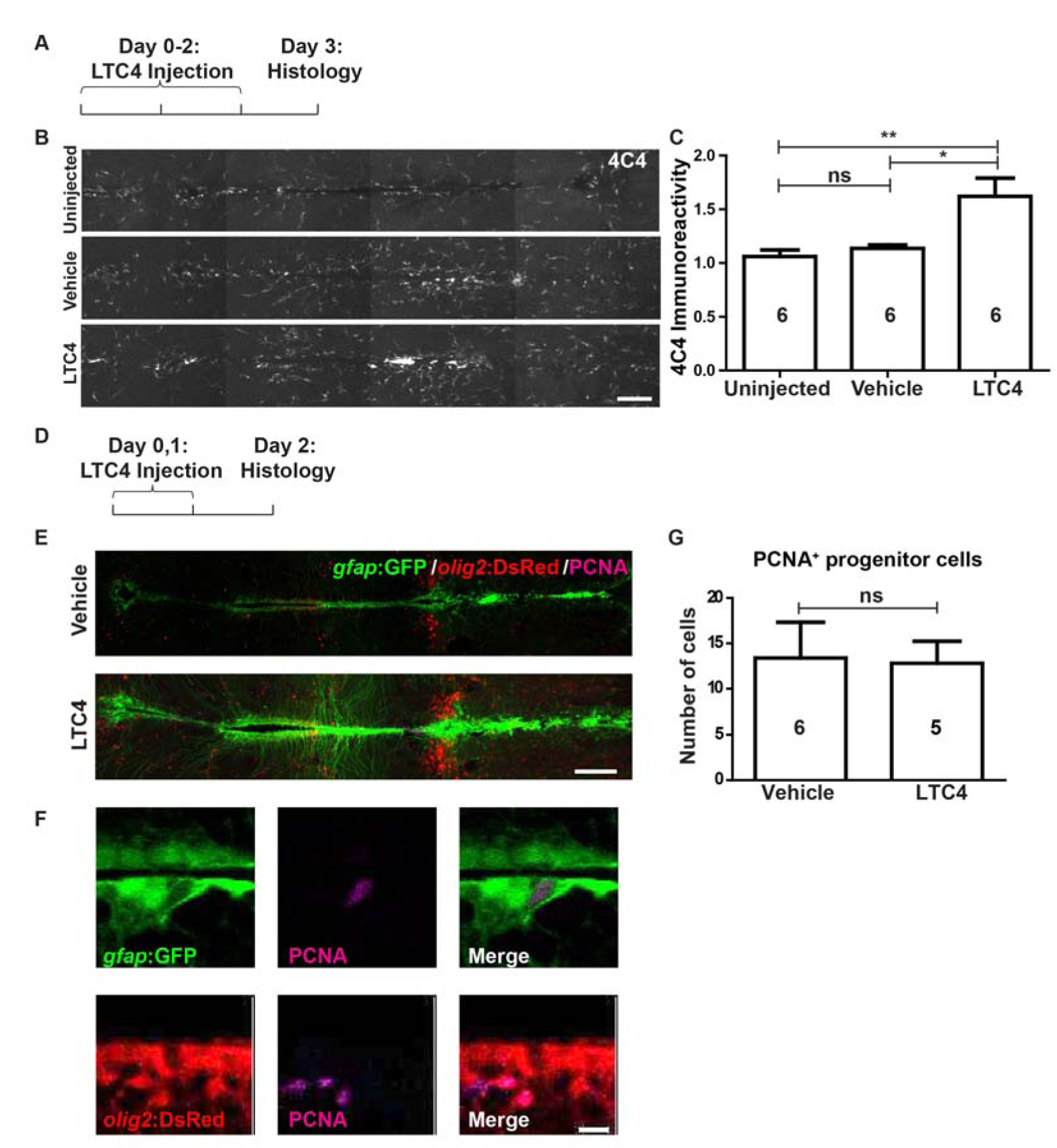
LTC4 moderately activates microglia but does not increase proliferation of ERGs. Horizontal sections are shown, rostral is left. **A-C:** LTC4, but not vehicle injection leads to an increase in microglia labelling in the brain **D-G:** PCNA labelling in *gfap*:GFP^+^ and/or *olig2:DsRed*^+^ ERGs is not increased by LTC4 (One-way ANOVA with Bonferroni post-hoc test used in C, Mann Whitney-U test used in G, *p < 0.05). Scale bars = 100 μm in B and E, 10 μm in F.

## REFERENCES

1 Jessberger, S. Neural repair in the adult brain. F1000Research 5, (2016).

2 Peron, S. & Berninger, B. Reawakening the sleeping beauty in the adult brain: neurogenesis from parenchymal glia. Curr Opin Genet Dev 34, 46-53, (2015).

3 Becker, C. G. & Becker, T. Neuronal regeneration from ependymo-radial glial cells: cook, little pot, cook! Dev Cell 32, 516-527, (2015).

4 Ghosh, S. & Hui, S. P. Regeneration of Zebrafish CNS: Adult Neurogenesis. Neural plasticity 2016, 5815439, (2016).

5 Alunni, A. & Bally-Cuif, L. A comparative view of regenerative neurogenesis in vertebrates. Development 143, 741-753, (2016).

6 Matsui, H. & Sugie, A. An optimized method for counting dopaminergic neurons in zebrafish. PLoS One 12, e0184363, (2017).

7 Vijayanathan, Y., Lim, F. T., Lim, S. M., Long, C. M., Tan, M. P., Majeed, A. B. A. & Ramasamy, K. 6-OHDA-Lesioned Adult Zebrafish as a Useful Parkinson’s Disease Model for Dopaminergic Neuroregeneration. Neurotoxicity research 32, 496-508, (2017).

8 Berg, D. A., Kirkham, M., Wang, H., Frisen, J. & Simon, A. Dopamine controls neurogenesis in the adult salamander midbrain in homeostasis and during regeneration of dopamine neurons. Cell Stem Cell 8, 426-433, (2011).

9 Tieu, K. A guide to neurotoxic animal models of Parkinson’s disease. Cold Spring Harbor perspectives in medicine 1, a009316, (2011).

10 Tay, T. L., Ronneberger, O., Ryu, S., Nitschke, R. & Driever, W. Comprehensive catecholaminergic projectome analysis reveals single-neuron integration of zebrafish ascending and descending dopaminergic systems. Nat Commun 2, 171, (2011).

11 Chen, Y. C., Priyadarshini, M. & Panula, P. Complementary developmental expression of the two tyrosine hydroxylase transcripts in zebrafish. Histochemistry and cell biology 132, 375-381, (2009).

12 McLean, D. L. & Fetcho, J. R. Relationship of tyrosine hydroxylase and serotonin immunoreactivity to sensorimotor circuitry in larval zebrafish. J Comp Neurol 480, 57-71, (2004).

13 McLean, D. L. &Fetcho, J. R. Ontogeny and innervation patterns of dopaminergic, noradrenergic, and serotonergic neurons in larval zebrafish. J Comp Neurol 480, 38-56, (2004).

14 Kuscha, V., Barreiro-Iglesias, A., Becker, C. G. & Becker, T. Plasticity of tyrosine hydroxylase and serotonergic systems in the regenerating spinal cord of adult zebrafish. J Comp Neurol 520, 933-951, (2012).

15 Lambert, A. M., Bonkowsky, J. L. & Masino, M. A. The conserved dopaminergic diencephalospinal tract mediates vertebrate locomotor development in zebrafish larvae. J. Neurosci. 32, 13488-13500, (2012).

16 Thirumalai, V. & Cline, H. T. Endogenous dopamine suppresses initiation of swimming in pre-feeding zebrafish larvae. J Neurophysiol, (2008).

17 Jay, M., De Faveri, F. &McDearmid, J. R. Firing dynamics and modulatory actions of supraspinal dopaminergic neurons during zebrafish locomotor behavior. Curr Biol 25, 435-444, (2015).

18 Reimer, M. M. et al. Dopamine from the Brain Promotes Spinal Motor Neuron Generation during Development and Adult Regeneration. Dev Cell 25, 478-491, (2013).

19 Wang, Y., Li, S., Liu, W., Wang, F., Hu, L. F., Zhong, Z. M., Wang, H. & Liu, C. F. Vesicular monoamine transporter 2 (Vmat2) knockdown elicits anxiety-like behavior in zebrafish. Biochem Biophys Res Commun 470, 792-797, (2016).

20 Tran, S., Nowicki, M., Muraleetharan, A., Chatterjee, D. & Gerlai, R. Neurochemical factors underlying individual differences in locomotor activity and anxiety-like behavioral responses in zebrafish. Prog Neuropsychopharmacol Biol Psychiatry 65, 25-33, (2016).

21 Grandel, H., Kaslin, J., Ganz, J., Wenzel, I. & Brand, M. Neural stem cells and neurogenesis in the adult zebrafish brain: origin, proliferation dynamics, migration and cell fate. Dev Biol 295, 263-277, (2006).

22 Grandel, H. & Brand, M. Comparative aspects of adult neural stem cell activity in vertebrates. Dev. Genes Evol. 223, 131-147, (2013).

23 Kyritsis, N., Kizil, C., Zocher, S., Kroehne, V., Kaslin, J., Freudenreich, D., Iltzsche, A. &Brand, M. Acute inflammation initiates the regenerative response in the adult zebrafish brain. Science 338, 13531356, (2012).

24 Ohnmacht, J., Yang, Y. J., Maurer, G. W., Barreiro-Iglesias, A., Tsarouchas, T. M., Wehner, D., Sieger, D., Becker, C. G. & Becker, T. Spinal motor neurons are regenerated after mechanical lesion and genetic ablation in larval zebrafish. Development, (2016).

25 Westerfield, M. The zebrafish book: a guide for the laboratory use of zebrafish (Danio rerio). 4th edn, (University of Oregon Press, 2000).

26 Kucenas, S., Takada, N., Park, H. C., Woodruff, E., Broadie, K. & Appel, B. CNS-derived glia ensheath peripheral nerves and mediate motor root development. Nat Neurosci 11, 143-151, (2008).

27 Bernardos, R. L. &Raymond, P. A. GFAP transgenic zebrafish. Gene Expr Patterns 6, 1007-1013, (2006).

28 Xi, Y., Yu, M., Godoy, R., Hatch, G., Poitras, L. & Ekker, M. Transgenic zebrafish expressing green fluorescent protein in dopaminergic neurons of the ventral diencephalon. Dev Dyn 240, 2539-2547, (2011).

29 Knopf, F., Schnabel, K., Haase, C., Pfeifer, K., Anastassiadis, K. & Weidinger, G. Dually inducible TetON systems for tissue-specific conditional gene expression in zebrafish. Proc. Natl. Acad. Sci. USA 107, 19933-19938, (2010).

30 Boniface, E. J., Lu, J., Victoroff, T., Zhu, M. & Chen, W. FlEx-based transgenic reporter lines for visualization of Cre and Flp activity in live zebrafish. Genesis 47, 484-491, (2009).

31 Ramachandran, R., Reifler, A., Parent, J. M. & Goldman, D. Conditional gene expression and lineage tracing of tuba1a expressing cells during zebrafish development and retina regeneration. J Comp Neurol 518, 4196-4212, (2010).

32 Skaggs, K., Goldman, D. & Parent, J. M. Excitotoxic brain injury in adult zebrafish stimulates neurogenesis and long-distance neuronal integration. Glia 62, 2061-2079, (2014).

33 Sallinen, V., Torkko, V., Sundvik, M., Reenila, I., Khrustalyov, D., Kaslin, J. & Panula, P. MPTP and MPP+ target specific aminergic cell populations in larval zebrafish. J Neurochem 108, 719-731, (2009).

34 Ohnmacht, J., Yang, Y., Maurer, G. W., Barreiro-Iglesias, A., Tsarouchas, T. M., Wehner, D., Sieger, D., Becker, C. G. & Becker, T. Spinal motor neurons are regenerated after mechanical lesion and genetic ablation in larval zebrafish. Development 143, 1464-1474, (2016).

35 Barreiro-Iglesias, A., Mysiak, K. S., Scott, A. L., Reimer, M. M., Yang, Y., Becker, C. G. & Becker, T. Serotonin Promotes Development and Regeneration of Spinal Motor Neurons in Zebrafish. Cell Rep 13, 924-932, (2015).

36 Reimer, M. M., Sorensen, I., Kuscha, V., Frank, R. E., Liu, C., Becker, C. G. & Becker, T. Motor neuron regeneration in adult zebrafish. J Neurosci 28, 8510-8516, (2008).

37 Becker, T. & Becker, C. G. Regenerating descending axons preferentially reroute to the gray matter in the presence of a general macrophage/microglial reaction caudal to a spinal transection in adult zebrafish. J. Comp. Neurol. 433, 131-147, (2001).

38 Kroehne, V., Freudenreich, D., Hans, S., Kaslin, J. & Brand, M. Regeneration of the adult zebrafish brain from neurogenic radial glia-type progenitors. Development 138, 4831-4841, (2011).

39 Blaser, R. & Gerlai, R. Behavioral phenotyping in zebrafish: comparison of three behavioral quantification methods. Behav Res Methods 38, 456-469, (2006).

40 Blaser, R. E. & Rosemberg, D. B. Measures of anxiety in zebrafish (Danio rerio): dissociation of black/white preference and novel tank test.PLoS One 7, e36931, (2012).

41 Stewart, A., Gaikwad, S., Kyzar, E., Green, J., Roth, A. & Kalueff, A. V. Modeling anxiety using adult zebrafish: a conceptual review. Neuropharmacology 62, 135-143, (2012).

42 Engeszer, R. E., Ryan, M. J. & Parichy, D. M. Learned social preference in zebrafish. Curr Biol 14, 881-884, (2004).

43 Pappas, S. S., Tiernan, C. T., Behrouz, B., Jordan, C. L., Breedlove, S. M., Goudreau, J. L. & Lookingland, K. J. Neonatal androgen-dependent sex differences in lumbar spinal cord dopamine concentrations and the number of A11 diencephalospinal dopamine neurons. J Comp Neurol 518, 2423-2436, (2010).

44 McPherson, A. D., Barrios, J. P., Luks-Morgan, S. J., Manfredi, J. P., Bonkowsky, J. L., Douglass, A. D. & Dorsky, R. I. Motor Behavior Mediated by Continuously Generated Dopaminergic Neurons in the Zebrafish Hypothalamus Recovers after Cell Ablation. Curr Biol 26, 263-269, (2016).

45 Godoy, R., Noble, S., Yoon, K., Anisman, H. & Ekker, M. Chemogenetic ablation of dopaminergic neurons leads to transient locomotor impairments in zebrafish larvae. J Neurochem 135, 249-260, (2015).

46 Kizil, C., Kyritsis, N., Dudczig, S., Kroehne, V., Freudenreich, D., Kaslin, J. & Brand, M. Regenerative neurogenesis from neural progenitor cells requires injury-induced expression of Gata3. Dev Cell 23, 1230-1237, (2012).

47 Kastenhuber, E., Kern, U., Bonkowsky, J. L., Chien, C. B., Driever, W. & Schweitzer, J. Netrin-DCC, Robo-Slit, and heparan sulfate proteoglycans coordinate lateral positioning of longitudinal dopaminergic diencephalospinal axons. J Neurosci 29, 8914-8926,(2009).

48 Wyatt, C., Ebert, A., Reimer, M. M., Rasband, K., Hardy, M., Chien, C. B., Becker, T. & Becker, C. G. Analysis of the astray/robo2 zebrafish mutant reveals that degenerating tracts do not provide strong guidance cues for regenerating optic axons. J. Neurosci. 30, 13838-13849, (2010).

49 Bhatt, D. H., Otto, S. J., Depoister, B. & Fetcho, J. R.Cyclic AMP-induced repair of zebrafish spinal circuits. Science 305, 254-258, (2004).

50 Becker, T., Bernhardt, R. R., Reinhard, E., Wullimann, M. F., Tongiorgi, E. & Schachner, M. Readiness of zebrafish brain neurons to regenerate a spinal axon correlates with differential expression of specific cell recognition molecules. J Neurosci 18, 5789-5803, (1998).

51 Anichtchik, O. V., Kaslin, J., Peitsaro, N., Scheinin, M. & Panula, P. Neurochemical and behavioural changes in zebrafish Danio rerio after systemic administration of 6-hydroxydopamine and 1-methyl-4-phenyl-1,2,3,6-tetrahydropyridine. J Neurochem 88, 443-453, (2004).

52 Lam, C. S., Korzh, V. & Strahle, U. Zebrafish embryos are susceptible to the dopaminergic neurotoxin MPTP. Eur J Neurosci 21, 1758-1762, (2005).

53 Bricaud, O., Chaar, V., Dambly-Chaudiere, C. & Ghysen, A. Early efferent innervation of the zebrafish lateral line. J Comp Neurol 434, 253-261., (2001).

54 Scerbina, T., Chatterjee, D. & Gerlai, R. Dopamine receptor antagonism disrupts social preference in zebrafish: a strain comparison study. Amino Acids 43, 2059-2072, (2012).

55 Kacprzak, V., Patel, N. A., Riley, E., Yu, L., Yeh, J. J. & Zhdanova, I. V. Dopaminergic control of anxiety in young and aged zebrafish. Pharmacology, biochemistry, and behavior 157, 1-8, (2017).

56 Parish, C. L., Beljajeva, A., Arenas, E. & Simon, A. Midbrain dopaminergic neurogenesis and behavioural recovery in a salamander lesion-induced regeneration model. Development 134, 2881-2887, (2007).

